# Esophageal epithelial *Ikkβ* deletion promotes eosinophilic esophagitis in experimental allergy mouse model

**DOI:** 10.1101/2024.07.05.602313

**Authors:** Margarette H Clevenger, Cenfu Wei, Adam L Karami, Lia E Tsikretsis, Dustin A Carlson, John E Pandolfino, Nirmala Gonsalves, Deborah R Winter, Kelly A Whelan, Marie-Pier Tétreault

## Abstract

**Background:** Eosinophilic esophagitis (EoE) is a chronic T helper type 2 (Th2)-associated inflammatory disorder triggered by food allergens, resulting in esophageal dysfunction through edema, fibrosis, and tissue remodeling. The role of epithelial remodeling in EoE pathogenesis is critical but not fully understood.

**Objective:** To investigate the role of epithelial IKKβ/NFκB signaling in EoE pathogenesis using a mouse model with conditional *Ikk*β knockout in esophageal epithelial cells (*Ikk*β*^EEC-KO^*).

**Methods:** EoE was induced in *Ikkβ^EEC-KO^*mice through skin sensitization with MC903/Ovalbumin (OVA) followed by intraesophageal OVA challenge. Histological and transcriptional analyses were performed to assess EoE features. Single-cell RNA sequencing (scRNA-seq) was used to profile esophageal mucosal cell populations and gene expression changes.

**Results:** *Ikkβ^EEC-KO^*/EoE mice exhibited hallmark EoE features, including eosinophil infiltration, intraepithelial eosinophils, microabscesses, basal cell hyperplasia, and lamina propria remodeling. RNA-seq revealed significant alterations in IKKβ/NFκB signaling pathways, with decreased expression of *RELA* and increased expression of IKKβ negative regulators. scRNA-seq analyses identified disrupted epithelial differentiation and barrier integrity, alongside increased type 2 immune responses and peptidase activity.

**Conclusion:** Our study demonstrates that loss of epithelial IKKβ signaling exacerbates EoE pathogenesis, highlighting the critical role of this pathway in maintaining epithelial homeostasis and preventing allergic inflammation. The *Ikkβ^EEC-KO^*/EoE mouse model closely mirrors human EoE, providing a valuable tool for investigating disease mechanisms and therapeutic targets. This model can facilitate the development of strategies to prevent chronic inflammation and tissue remodeling in EoE.

**Key Messages:**

- Critical Role of Epithelial IKKβ/NFκB Signaling: Loss of this signaling exacerbates EoE, causing eosinophil infiltration, basal cell hyperplasia, and tissue remodeling, highlighting its importance in esophageal health.
- Molecular Insights and Therapeutic Targets: scRNA-seq identified disrupted epithelial differentiation, barrier integrity, and enhanced type 2 immune responses, suggesting potential therapeutic targets for EoE.
- Relevance of the *Ikkβ^EEC-KO^*/EoE Mouse Model: This model replicates human EoE features, making it a valuable tool for studying EoE mechanisms and testing treatments, which can drive the development of effective therapies.

**Capsule Summary:** This study reveals the crucial role of epithelial IKKβ/NFκB signaling in EoE, providing insights into disease mechanisms and potential therapeutic targets, highly relevant for advancing clinical management of EoE.

## Introduction

Eosinophilic esophagitis (EoE) is a chronic Th2-associated inflammatory disorder triggered by food allergens, resulting in esophageal dysfunction through edema, fibrosis, and wall remodeling^14, 15, 16^. Despite treatment advancements, many patients experience recurrence or are unresponsive^1–3^, leading to reduced quality of life and high healthcare costs^4, 5^. Therefore, better molecular characterization of EoE is needed to develop effective therapies.

EoE is diagnosed by esophageal dysphagia and biopsies showing at least 15 eosinophils per high-power field. However, key epithelial changes such as dilation of intercellular spaces (DIS), loss of differentiation, and basal cell hyperplasia (BCH) are also known to drive the disease^6–9^. Supporting the important contribution of epithelial remodeling in disease pathogenesis, improved histological scoring systems focusing on these changes outperform peak eosinophil counts as diagnostic tools^10^. Furthermore, recent studies show EoE can be driven by epithelial-derived mediators, independent of the adaptive immune system^11^.

The IKKβ/NFκB pathway is a central hub for inflammatory responses, controlling processes like inflammation, immunity, cell survival, and growth^12–15^. Dysregulation of this pathway is linked to numerous inflammatory diseases and cancers^12–15^. However, most studies of esophageal diseases have focused on IKKβ or NFκB expression levels without functional *in vivo* analyses. Investigations in mice with conditional *Ikk*β overexpression or knockout in various tissues have demonstrated the context-dependent roles of this pathway^16–21^, highlighting the importance of studying it in different tissues and disease states.

This study examines the role of epithelial IKKβ in EoE pathogenesis by using mice with conditional *Ikk*β knockout in esophageal epithelial cells (EEC), subjected to skin sensitization with MC903/Ovalbumin (OVA) followed by intraesophageal OVA challenge (*Ikkβ^EEC-KO^*/EoE). We show that *Ikkβ^EEC-KO^*/EoE mice replicate key histological and transcriptional changes observed in human EoE, including eosinophil infiltration, intraepithelial eosinophilia, microabscesses, DIS, BCH, CD4+ T cell recruitment, and lamina propria remodeling, demonstrating the critical role of epithelial IKKβ in EoE pathogenesis.

## Materials and Methods

### Generation of ED-L2-Cre; Ikkβ^L/L^ Mice

All animal studies were approved by the IACUC at Northwestern University. To generate *ED-L2- Cre; Ikkβ^L/L^* mice*, floxed Ikkβ^L/L^* mice^22^ were crossed with *EBV-ED-L2/Cre* mice^23^ (Balb/C). For experiments, sex-matched littermate *Ikkβ^L/L^* mice (control) served as controls. Both male and female groups were included. Additional details are in the Supplementary Materials and Methods section.

### Induction of EoE-like disease in mice

EoE-like disease was induced in three-month old mice using a previously described methodology ^24^. Mice were treated topically on the ears with MC903 (2nmol) and OVA (100 μg) once daily for 12 days. On days 15 and 17, mice were challenged intraesophageally (IE) with 50 mg OVA. On day 15, mice were given access to OVA water (1.5g/L) ad libitum until sacrifice (day 18). Additional details are in **Supplemental Table 1** and in the Supplementary Materials and Methods section.

### Immunohistochemistry, Immunofluorescence and scoring

Immunostaining was performed using standard protocols. For protocols, histological scoring, staining quantification and antibodies used, see **Supplemental Table 2** and the Supplementary Materials and Methods section.

### Western Blots

Western Blot were performed as previously described^25^. More information can be found in the Supplementary Materials and Methods section.

### Human specimen sample collection and processing

Details regarding specimen collection, inclusion and exclusion criteria can be found in the Supplementary Materials and Methods section and in **Supplemental Table 3**. All procedures using human tissue were approved by the Northwestern Institutional Review Board (STU00208111) and conducted according to the relevant guidelines and regulations. Informed consent was obtained from all subjects/legal guardians prior to participation.

### Bulk RNA sequencing

RNA extraction, DNA libraries generation, sequencing, data filtering and analysis were performed as previously described^26^. Additional details can be found in the Supplementary Materials and Methods section. RNA-seq data was deposited in Gene Expression Omnibus (GEO #270219, #271128) and can be accessed at http://www.ncbi.nlm.nih.gov/geo/query/acc.cgi?acc=GSE270219, http://www.ncbi.nlm.nih.gov/geo/query/acc.cgi?acc=GSE271128.

### ScRNA-seq

Single cell suspensions of mouse esophageal mucosal cells were generated. Library preparation, sequencing and computational analyses were performed as detailed in the Supplementary Materials and Methods section.

### Data and code availability

Raw sequencing files and processed data are deposited in NCBI’s Gene Expression Omnibus (GEO) database under accession codes GSE270219 and GSE271128. Analytic code is available at the ’scRNA-Mouse_EoE_Esophagus’ repository hosted by the Tetreault Lab on GitHub.

### Statistical Analyses

Statistical analyses were conducted using R version 4.1.1. Descriptive statistics are presented as mean ± standard error of the mean for continuous variables and frequency counts for categorical variables. Additional details can be found in the Supplementary Materials and Methods section.

## Results

### Decreased IKKβ/NFκB signaling is detected in both human EoE and in a mouse model of EoE-like inflammation

Recent RNA-seq analysis of esophageal mucosal biopsies from 19 adult EoE patients and 8 healthy controls (HC) revealed aberrant gene expression in EoE. Gene set enrichment analysis (GSEA) demonstrated an enrichment of IKKβ/NFκB signaling in both proximal (**Figure 1A**) and distal (**Figure 1B**) esophagus. Differentially expressed genes revealed an up-regulation of several negative regulators of IKKβ/NFκB signaling and a down- regulation of positive regulators, including a significant decrease in *RELA* expression and an increase in *IKBIP* expression (**Figure 1C-D**). To assess IKKβ/p65 NFκB signaling activity in mouse EoE, we induced EoE-like inflammation through ear sensitization of Balb/c mice with the vitamin D analog MC903 and Ovalbumin (OVA) followed by intra-esophageal (IE) gavage with OVA (**Figure 1E**)^24^. As shown in **Figure 1F**, decreased phosphorylation of p65 NFκB was observed in mice with EoE-like inflammation compared to controls, suggesting that decreased epithelial IKKβ/NFκB signaling contributes to EoE pathogenesis.

**Figure 1.**
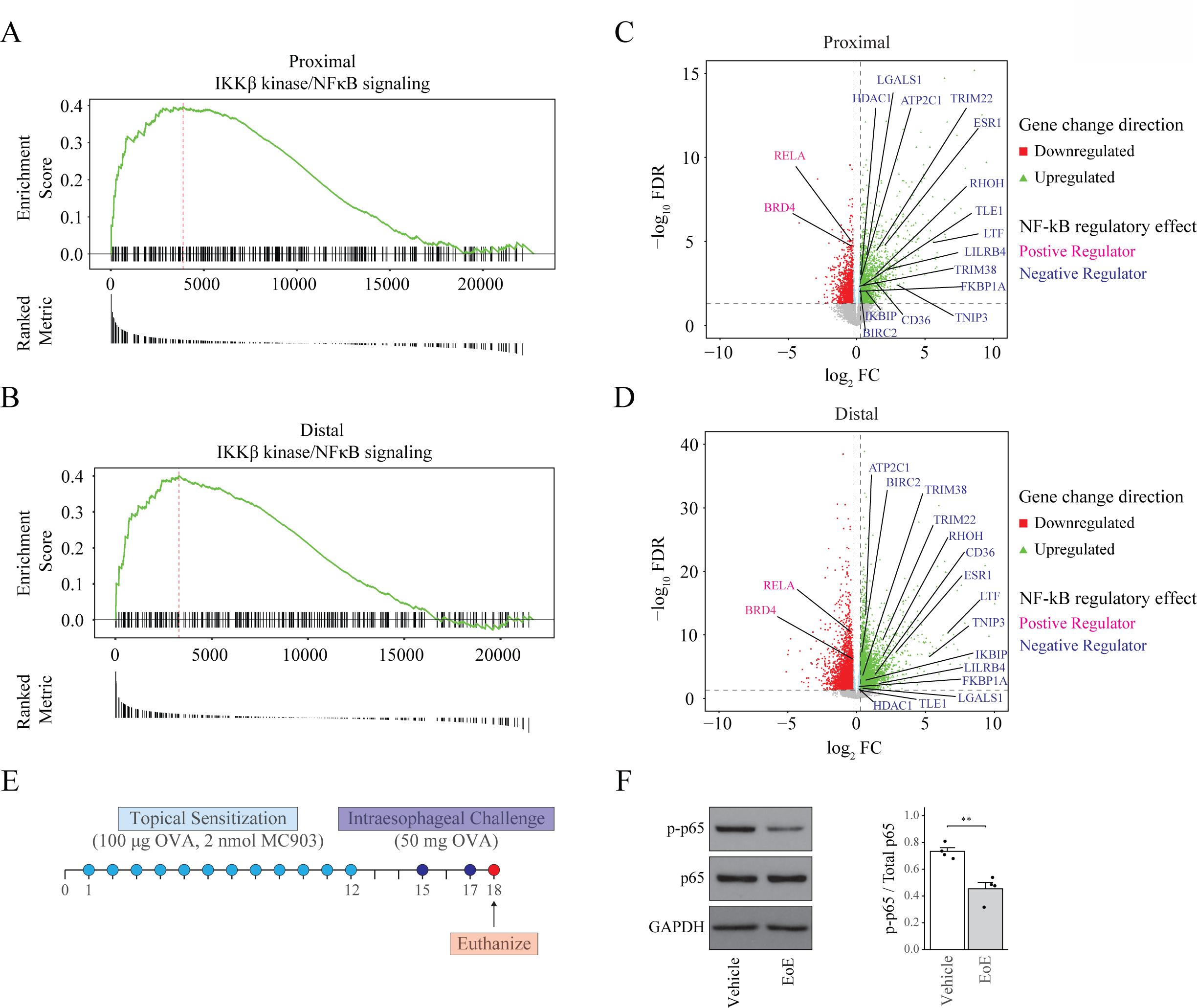
Reduced canonical NF-κB signaling is detected in the esophageal mucosa of human and mouse EoE. (**A, C**) Running enrichment score plots show the enrichment score curve (green) and gene members from the ranked list (EoE vs. HC) in proximal (**A**) and distal (**C**) bulk RNA sequencing datasets, highlighting IKKβ/NF-κB signaling genes identified by GSEA. Red line = point of maximum enrichment score for the gene set; black lines = IKKβ/NF- κB signaling members appearing in the ranked gene list. The ranked metric shows the ranking metric (log2FC) value for each gene. (**B, D**) Volcano plots displaying log2FC vs. -log10(FDR) of DEGs from EoE vs. HC differential expression analysis in proximal (**B**) and distal (**D**) datasets. Dashed lines indicate |log2FC)| = 0.26 (vertical); -log10(FDR) = 1.3 (horizontal). Downregulated DEGs = red squares, upregulated DEGs = green triangles. Positive NF-κB regulators = pink, negative NF-κB regulators = blue. (**E**) Schematic of MC903/OVA EoE treatment. (**F**) Left: Western blot showing p65 and p65^ser536^ expression in EoE mice, with GAPDH as a loading control. Right: Densitometric analysis of the ratio of phosphorylated to total p65 protein ratio. n= 4 mice. 2-tailed Student t-test, **P ≤ 0.01.

### Intraesophageal challenge of OVA in epicutaneously sensitized mice with loss of epithelial Ikkβ results in EoE

To investigate the role of decreased epithelial IKKβ/NFκB signaling in EoE pathogenesis, we generated mice with conditional esophageal epithelial *Ikk*β deletion (*Ikkβ^EEC-KO^*) by crossing *Ikk*β floxed mice (*Ikk*β*^L/L^*, controls)^27^ with *ED-L2/Cre* mice^23^, and then backcrossed onto the Balb/c background. *Ikkβ^EEC-KO^* mice up to 6 months-old were phenotypically normal (**Figure 2A, Supplemental Figure 1A**). For EoE induction, both *Ikkβ^EEC-KO^*mice (*Ikkβ^EEC-KO^*/EoE) and their littermate controls (control/EoE) were treated with MC903/OVA followed by intraesophageal OVA challenge (**Figure 1E)**. Experimental groups also included untreated *Ikk*β*^ECC-KO^*mice and their littermate controls, epicutaneously sensitized mice without intraesophageal challenge, and vehicle-gavaged mice following sensitization. No abnormal findings were observed in these groups (**Supplemental Figure 1A**).

**Figure 2.**
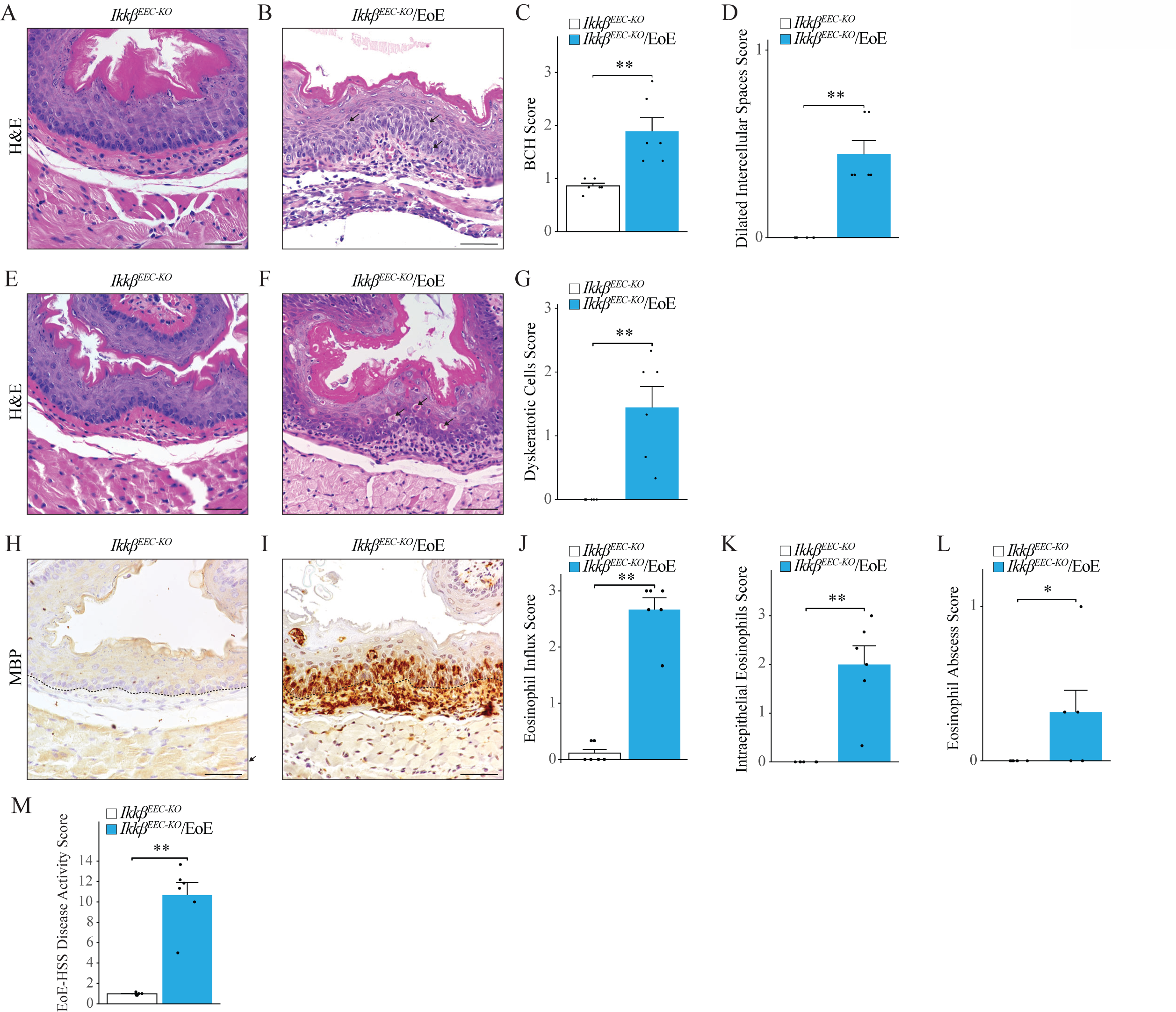
Experimental EoE with esophageal epithelial *Ikkβ* loss recapitulates histological features of human disease. (**A-D**) H&E staining of esophageal sections of *Ikkβ^EEC-KO^* (**A**) and *Ikkβ^EEC-KO^*/EoE mice (**B**) (BCH area = dashed black line, DIS = black arrows). (**C-D**) Quantification of BCH (**C**) and DIS (**D**) scoring. (**E-F**) H&E staining of esophageal sections of *Ikkβ^EEC-KO^* (**E**) and *Ikkβ^EEC-KO^*/EoE mice (**F**) with dyskeratotic cells (black arrows), quantified in (**G**). (**H-I**) MBP immunohistochemistry of esophageal sections of *Ikkβ^EEC-KO^* (**H**) and *Ikkβ^EEC-KO^*/EoE (**I**) mice. Dashed black lines = basement membrane. (**J**) Total eosinophil influx, intraepithelial eosinophil influx (**K**) and eosinophil abscess (**L**) scoring. (**M**) EoE-HSS disease activity scoring. Scale bar: 50 µm. n= 6 mice. Bar graphs represent means ± SEM. 2-tailed Student t-test, \**P* ≤ 0.05, \*\**P* ≤ 0.01.

Unlike control mice, which showed minimal eosinophil recruitment and no intraepithelial eosinophils with MC903/OVA treatment, as previously reported^24^ (**Figure 2A**), *Ikkβ^EEC-KO^*/EoE mice developed key EoE histological features, including BCH, DIS (**Figure 2B-D**), dyskeratotic cells (**Figure 2E-G**), extensive eosinophilic infiltration, intraepithelial eosinophilia and microabscesses (**Figure 2H-L, Supplemental Figure 1A, 2**). Disease activity, assessed using EoE-HSS criteria, was significantly higher disease activity in *Ikkβ^EEC-KO^*/EoE mice compared to all other groups (**Figure 2M, Supplemental Figure 1B**). Eosinophilic infiltration, assessed via anti-MBP staining, showed no eosinophils in untreated, vehicle gavaged or control groups (**Supplemental Figure 2**). Minor eosinophil presence was observed in control/EoE, and sensitized mice, but less pronounced than in *Ikkβ^EEC-KO^*/EoE mice (**Supplemental Figure 2**). Importantly, only *Ikkβ^EEC-KO^*/EoE mice met EoE diagnostic criteria. Masson Trichrome staining showed increased collagen remodeling in *Ikkβ^EEC-KO^*/EoE mice compared to all other groups (**Supplemental Figure 3**), indicating that *Ikkβ^EEC-KO^*/EoE mice display the comprehensive histological characteristics of human EoE.

### Identification of esophageal mucosal cell populations in mouse EoE

To identify the target genes permitting EoE development downstream of epithelial *Ikkβ* loss and evaluate transcriptional similarity to human EoE, we performed scRNA-seq on esophageal mucosa from *Ikkβ^EEC-KO^*/EoE mice, control mice, and relevant control groups. Single-cell suspensions were generated from the esophageal mucosal tissue and sequencing was performed using the 10x Genomics platform. After quality control, samples were integrated using reciprocal principal component analysis (rPCA) for dimensional reduction with the Seurat R package. Uniform Manifold Approximation and Projection for Dimension Reduction (UMAP) and unsupervised graph-based clustering then identified esophageal mucosal cell types using established markers (**Figure 3A-B**).

**Figure 3.**
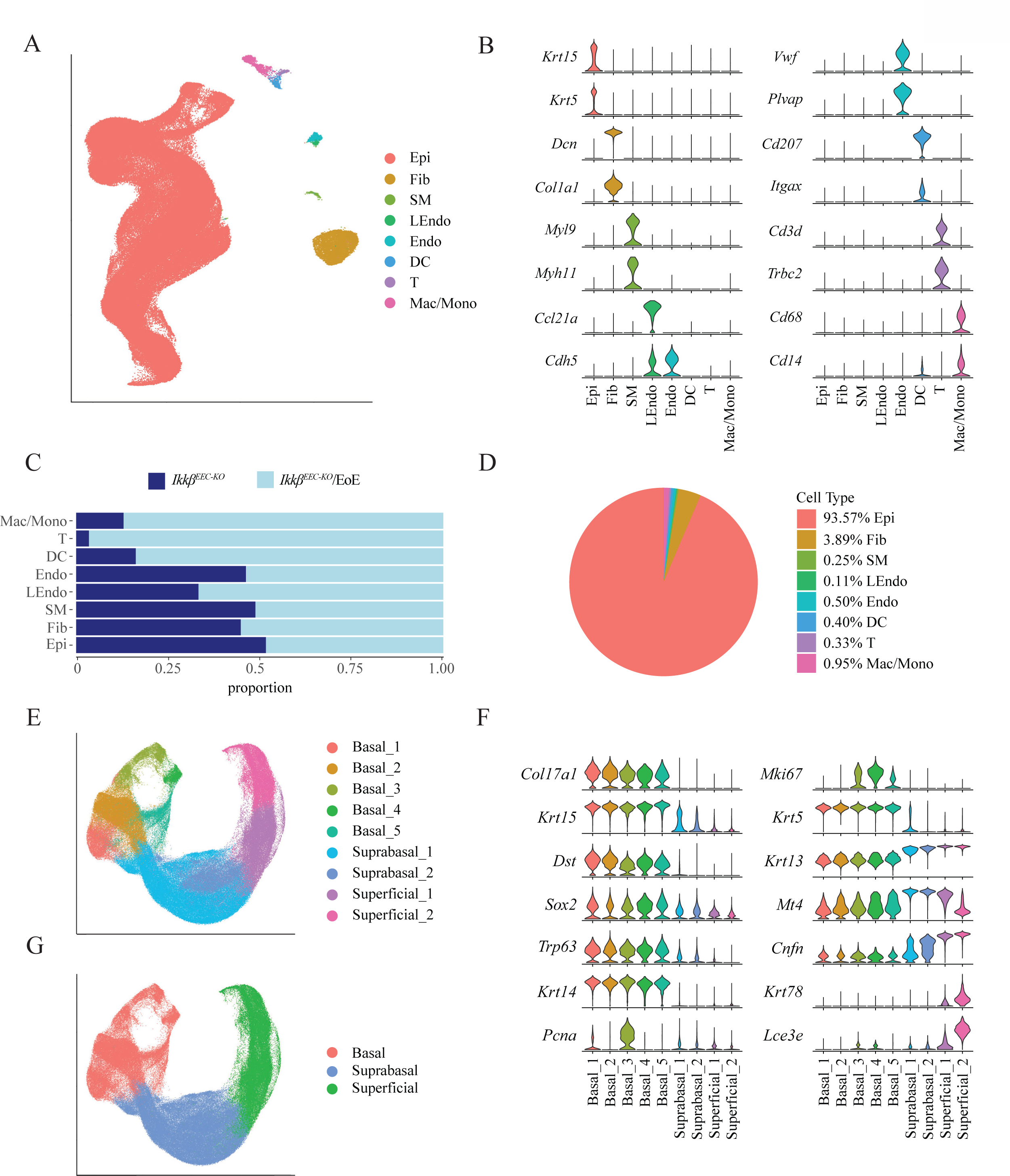
Single-cell transcriptomic landscape of esophageal mucosal cells in the murine EoE dataset. (**A**) UMAP visualization of unsupervised clustering of mouse esophageal mucosal cells by cell type. (**B**) Violin plots of cell-type specific gene marker expression. (**C**) Bar plot showing the frequency of each esophageal mucosal cell type in *Ikkβ^EEC-KO^*and *Ikkβ^EEC-KO^*/EoE mice. (**D**) Pie chart showing the proportion of each cell type in the murine EoE dataset. (**E**) UMAP visualization of epithelial sub-clustering (colors = EEC cluster). (**F**) Violin plots of marker gene expression across EEC clusters. (**G**) UMAP visualization of the EEC dataset (colors = epithelial compartments).

Our integrated dataset encompassed a total of 203,476 cells, annotated into eight major cell types: epithelial cells (Epi) (n = 190,389), fibroblast cells (Fib) (n = 7,919), smooth muscle cells (SM) (n = 511), lymphatic endothelial cells (LEndo) (n = 232), endothelial cells (Endo) (n = 1013), dendritic cells (DC) (n = 817), T cells (T) (n = 669) and macrophage and monocyte cells (Mac/Mono) (n = 1925) (**Figure 3A**). Cell type annotations were validated through inter-cluster transcriptional profiles comparisons (**Supplemental Figure 4A**). The cell population distribution was similar between *Ikk*β*^ECC-KO^/EoE* and *Ikk*β*^ECC-KO^*mice, except for differences in lymphatic endothelial cells, dendritic cells, T cells and macrophages/monocytes (**Figure 3C, Supplemental Figure 4B-C**). EECs was the dominant cell type (**Figure 3D**).

### Defining clusters of EECs

We next investigated the transcriptional changes occurring in *Ikkβ^EEC-KO^*/EoE EEC. Using unsupervised graph-based clustering, we identified six epithelial clusters (**Supplemental Figure 5A**), which were mapped to epithelial compartments (basal (B), suprabasal (SB), superficial (SF)) using expression markers from our untreated mouse dataset (**Supplemental Figure 5B**)^28, 29^. Basal clusters were further subclustered into quiescent and cell cycle phases^28^, resulting in 5 basal subclusters (B1-B5) (**Supplemental Figure 5C-E**), raising the total to 9 distinct epithelial clusters (**Figure 3E**). In the basal compartment, clusters B1 and B2 represented quiescent cells, marked by high expression of *Col17a1*, *Krt15*, and *Dst*. Clusters B3-B5 comprised actively proliferating EEC, with B3 showing increased levels of the S-phase marker *Pcna*, B4 showing high expression of the G2/M marker *Mki67*, and B5 indicating a transition out of the cell cycle with lower *Mki67* levels, (**Figure 3F**). Suprabasal clusters were marked by *Krt13* and *Mt4* expression, while superficial clusters were distinguished by early and late differentiation markers: *Cnfn* for early SF1 and *Krt78* and *Lce3e* for late SF2 clusters (**Figure 3F**). Cluster composition analysis in biological replicates revealed a degree of representation for each cluster across each mouse group (**Supplemental Figure 5F**), with consistent clustering across all experimental conditions (**Supplemental Figure 5G**). These 9 epithelial clusters were mapped to epithelial compartments (**Figure 3G**), and cluster annotation was validated using the transcriptional profile of each untreated *IKK*β*^ECC-KO^*epithelial cell cluster (**Supplemental Figure 6A**). A detailed schematic of the different epithelial clusters and compartments, with the corresponding markers is illustrated in **Supplemental Figure 6B**.

### ScRNA-seq highlights physiological similarities between *Ikkβ^EEC-KO^*/EoE mice and human EoE

To assess the physiological relevance of *Ikkβ^EEC-KO^*/EoE mice to human EoE, we performed differential gene expression analysis across all mouse group and compared the differentially expressed genes (DEGs) with those previously established in human EoE^28^. Our analysis revealed that loss of epithelial *Ikkβ,* the mechanical impact of gavage or sensitization alone resulted in few significant EoE-related DEGs, mirroring the patterns observed in control/EoE mice (**Figure 4A**). However, a notable overlap of over 200 DEGs was observed when comparing *Ikkβ^EEC-KO^*/EoE mice to *Ikkβ^EEC-KO^* mice, and approximately 100 DEGs aligned with human EoE when comparing *Ikkβ^EEC-KO^*/EoE mice and control/EoE mice (**Figure 4A**). These findings suggest the physiological relevance of *Ikkβ^EEC-KO^*/EoE mice for studying EoE pathogenesis.

**Figure 4.**
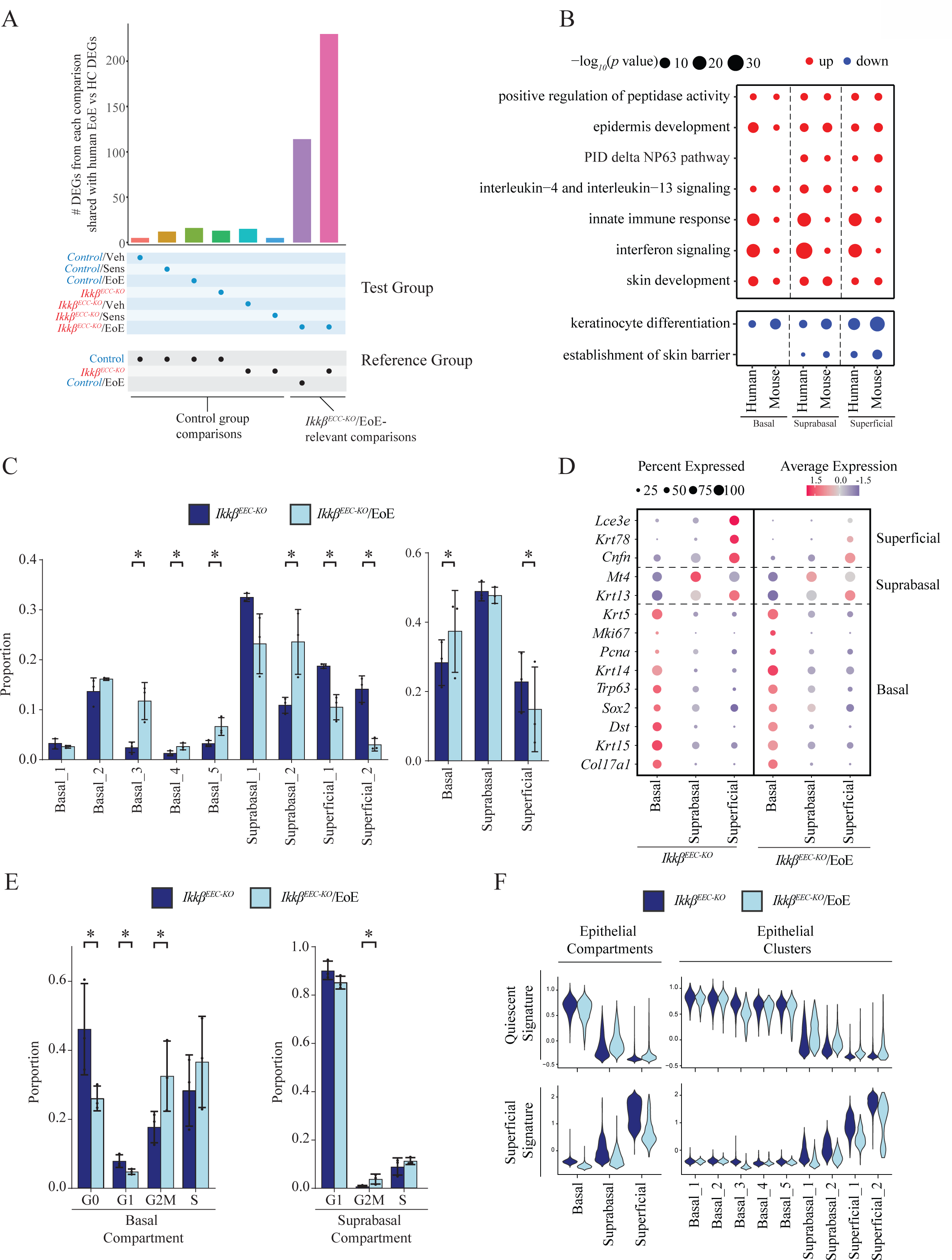
Transcriptomic changes in gene signaling and cellular identities in *Ikkβ^EEC-KO^*/EoE mice compared to human EoE. (**A**) Bar plot showing DEGs count overlapping between murine EoE dataset group comparisons and human EoE vs HC scRNA-seq dataset. FDR < 0.05, logFC > 0.25. Tested group and reference groups are indicated below the plot. (**B**) Dot plot of enriched pathways from DEGs in human EoE versus HC or *Ikkβ^EEC-KO^*/EoE versus *Ikkβ^EEC-KO^* mice. (**C**) Bar plots showing the cell proportions in each EEC cluster or compartment in *Ikkβ^EEC-KO^* or *Ikkβ^EEC-KO^*/EoE mice as a fraction of total EEC per treatment group. Statistics by permutation testing, n = 10,000. (**D**) Average expression z-scores of basal and differentiated EEC gene markers in *Ikkβ^EEC-KO^*and *Ikkβ^EEC-KO^*/EoE mice across epithelial compartments. Dot size indicates cell percentages per compartment with gene expression. Color gradient = average expression level. (**E**) Bar plots showing cell proportions in each cell cycle stage in basal or suprabasal EEC compartments in *IKKβ^EEC-KO^* and *IKKβ^EEC-KO^*/EoE mice as a fraction of total EEC per compartment. Statistics by permutation testing, n = 10,000. (**F**) Violin plots of quiescent or superficial gene signatures scoring in EEC compartments or clusters between *Ikkβ^EEC-KO^* and *Ikkβ^EEC-KO^*/EoE mice.

Our control group analysis showed that *Ikk*β deletion, sensitization alone, or intraesophageal gavage had minimal effects on EoE development. Therefore, we focused further analyses on comparing *Ikkβ^EEC-KO^*/EoE with untreated *Ikkβ^EEC-KO^* mice. Differential gene expression and pathway enrichment analysis comparing the epithelial compartments of *Ikkβ^EEC-KO^*/EoE mice with those of untreated *Ikkβ^EEC-KO^* mice revealed that, similar to human EoE, *Ikkβ^EEC-KO^*/EoE mice showed activation in pathways including IL-4 and IL-13 signaling, positive regulation of peptidase activity, skin and epidermis development, along with innate immune and interferon signaling (**Figure 4B**). Conversely, other EoE-associated pathways such as keratinocyte differentiation and skin barrier formation were down-regulated (**Figure 4B**). Increased CD4+ T lymphocyte recruitment in *Ikkβ^EEC-KO^*/EoE mice further supported the activation of IL-13/IL-4 signaling (**Supplemental Figure 7A-B**). These findings suggest that *Ikkβ^EEC-KO^*/EoE mice mirror changes seen in human EoE, indicating that the allergic response developed through epicutaneous MC903/OVA sensitization and intraesophageal OVA gavage is contingent on the absence of *Ikk*β in some EEC.

### Decreased terminal EEC differentiation and increased proliferation in Ikkβ^EEC-KO^/EoE mice

We next investigated changes in cellular identity by analyzing EEC distribution across epithelial clusters and compartments in *Ikkβ^EEC-KO^*/EoE mice. Compared to *Ikkβ^EEC-KO^* mice, *Ikkβ^EEC-KO^*/EoE mice showed a decrease in superficial cells, with a shift from SB1 to SB2 clusters (**Figure 4C, Supplemental Figure 8A**). There was also an increase in actively cycling basal clusters B3-B5 (**Figure 4C, Supplemental Figure 8B**). We then analyzed differentiation marker expression in EEC from *Ikkβ^EEC-KO^*/EoE mice, finding downregulation of the critical terminal differentiation and cornification markers *Lce3e* and *Krt78* across suprabasal and superficial layers, a reduction of the percentage of cells expressing these genes, and an abnormal expansion of *Sox2* expression in the suprabasal layer (**Figure 4D**). The early differentiation genes *Krt13* and *Mt4* were upregulated in the suprabasal layer (**Figure 4D**), but less so than in untreated *Ikkβ^EEC-KO^* mice. To confirm increased proliferation in the basal and suprabasal compartments in EoE, we examined G2/M cell division rates using published signatures^30, 31^, finding a significant increase in the basal compartment (**Figure 4E, Supplemental Figure 8B**). Ki-67 staining confirmed these findings (**Supplemental Figure 9**). Following our work in human EoE^28^, we developed gene signatures to identify genes preferentially expressed in quiescent (B1-B2) or superficial (SF1-SF2) EEC in untreated mice (**Supplemental Table 4**). *Ikkβ^EEC-KO^*/EoE mice showed a significant decrease in the superficial score in the suprabasal and superficial compartments, with an increase in quiescent gene expression (**Figure 4F)**, indicating altered differentiation and maintained basal identity in these compartments.

Using Monocle3 for pseudotemporal analysis on merged epithelial samples, as we have previously done (**Figure 5A**)^28, 29^, we observed that EECs from *Ikkβ^EEC-KO^* mice followed the standard differentiation pathway (**Figure 5B-D**). In contrast, *Ikkβ^EEC-KO^*/EoE mice showed a divergence during the intermediate differentiation stage, with more cells at this stage and fewer progressing to the late superficial stage (**Figure 5B-D**). This shift was consistently observed across all biological replicates (**Supplemental Figure 10**).

**Figure 5.**
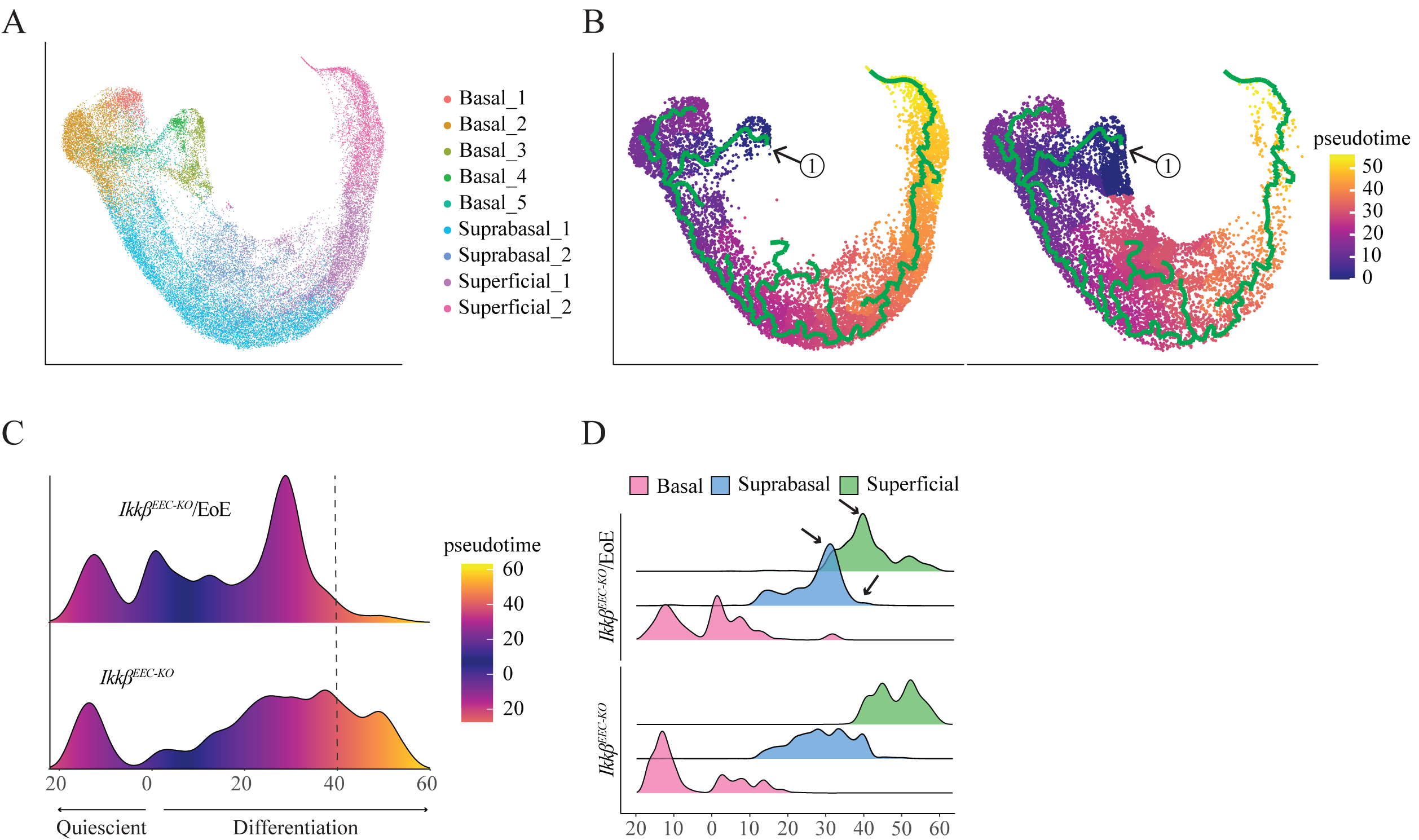
Pseudotime trajectory analysis of EEC in *Ikkβ^EEC-KO^*and *Ikkβ^EEC-KO^*/EoE mice. (**A**) UMAP of merged *Ikkβ^EEC-KO^* and *Ikkβ^EEC-KO^*/EoE dataset, colored by integrated EEC clusters. (**B**) Pseudotime trajectory analysis of merged *Ikkβ^EEC-KO^* and *Ikkβ^EEC-KO^*/EoE EEC dataset. The trajectory (green line) starts from cells in S-phase (Basal_3) and progresses towards quiescence or terminal differentiation. (**C**) Ridge plot showing pseudotime value distribution across all EEC from *Ikkβ^EEC-KO^*or *Ikkβ^EEC-KO^*/EoE mice. (**D**) Ridge plot showing the pseudotime value distribution in each EEC compartment from *Ikkβ^EEC-KO^*or *Ikkβ^EEC-KO^*/EoE mice. Arrows = areas/differential density peaks between the two groups.

### Role of epithelial Ikkβ in modulating EoE pathology

Wild-type mice subjected to MC903/OVA sensitization/intraesophageal OVA gavage show a mild inflammatory response that does not meet EoE diagnostic criteria. However, mice with conditional loss of esophageal epithelial *Ikkβ* (*Ikkβ^EEC-KO^*/EoE) display hallmark pathological characteristics of EoE (**Figure 2**). These molecular features mirror human EoE pathology, prompting further investigation of genes impacted by the absence of esophageal epithelial *Ikk*β. We conducted differential gene expression analysis of EEC from basal, suprabasal and superficial compartments in *Ikkβ^EEC-KO^*/EoE mice compared to control/EoE mice. Analysis revealed 440 shared DEGs among the 1,027 altered genes in *Ikkβ^EEC-KO^*/EoE mice, indicating that these genes are associated with the absence of epithelial *Ikkβ* (**Figure 6A**). Pathway enrichment analysis of these 440 genes highlighted key EoE-associated pathways, particularly late-stage differentiation processes, cornified envelope formation and barrier function establishment (**Figure 6B**). This suggests that loss of epithelial *Ikkβ* primarily impacts differentiation and barrier integrity, potentially acting as a precursor to disease progression. Other impacted pathways included regulation of type 2 immune responses, antigen processing and presentation, and endopeptidase activity (**Figure 6B**).

**Figure 6.**
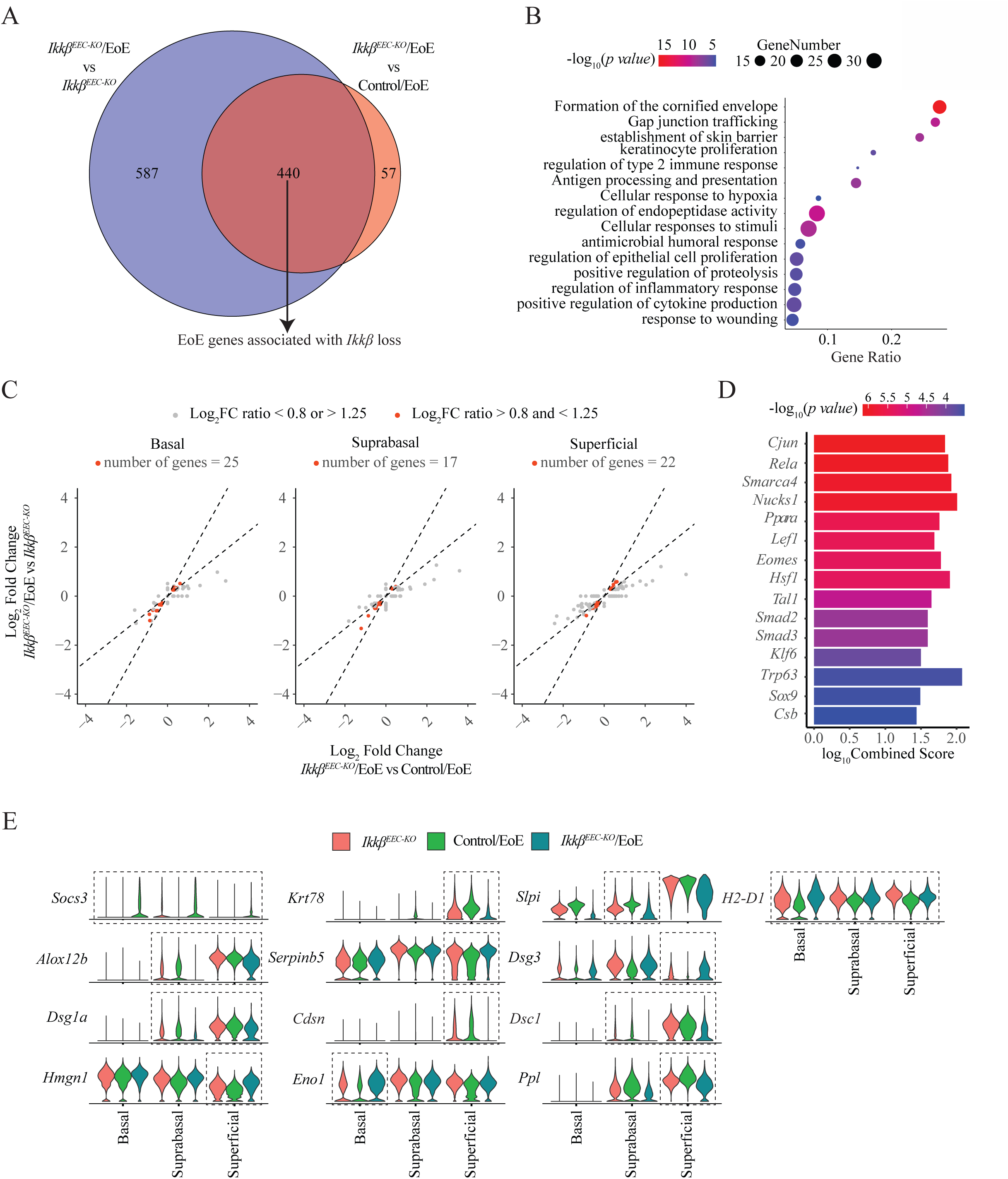
Identification of EoE-related genes associated with loss of esophageal epithelial *Ikk*β. (**A**) Venn diagram showing overlap of DEGs between *Ikkβ^EEC-KO^*/EoE vs *Ikkβ^EEC-KO^* and vs control/EoE mice to identify DEGs linked to *Ikk*β loss in EoE. (**B**) Dot plot of enriched pathways from the 440 overlapping DEGs identified in (**A**). (**C**) Scatter plots of log2FC for the 440 overlapping DEGs from *Ikkβ^EEC-KO^*/EoE vs *Ikkβ^EEC-KO^* (y-axis) or vs *Ikkβ^EEC-KO^* (x-axis) in each EEC compartment. Dashed lines with slopes of 0.8 and 1.25 indicate selection criteria references for DEGs with similar expression trends, highlighting 62 DEGs in red. (**D**) Bar plot of predicted upstream transcription factors for the DEGs from (**C**). (**E**) Violin plot showing expression levels of selected genes from (**C**) across EEC compartments in *Ikkβ^EEC-KO^*, control/EoE and *Ikkβ^EEC-KO^*/EoE mice.

We next examined the expression profiles of the 440 DEGs linked to EoE pathways and specifically attributable to epithelial *Ikk*β loss. We aimed to identify genes with consistent changes across conditions, indicating their regulation by IKKβ and their relevance to disease progression. Expression patterns were examined across different epithelial compartments in *Ikkβ^EEC-KO^*, *Ikkβ^EEC-KO^*/EoE, and control/EoE mice (**Supplemental Figure 11).** We evaluated the log fold change (logFC) values of these DEGs, comparing *Ikkβ^EEC-KO^*/EoE mice to *Ikkβ^EEC-KO^* mice (y-axis) and control/EoE mice (x-axis) for each epithelial compartment (**Figure 6C**). Genes with logFC ratios between 0.8-1.25 were selected, narrowing the analysis to 64 genes (**Figure 6C**). EnrichR analysis^32, 33^ identified *Rela*, encoding the NFκB p65 subunit, as a top-predicted transcription factor, potentially regulating 23% of these genes (**Figure 6D**). The expression of a subset of these 64 genes across epithelial cell compartments is shown in **Figure 6E**. Key genes regulated by *Ikkβ* in EoE development include decreased expression of the desmosomal cadherin protein *Dsg1a*^34^ and the intermediate filament keratin *Krt78*^35^, associated with loss of barrier function and differentiation. Other notable genes include loss of *Ppl,* involved in desmosome linkage to intermediate filaments and *Cdsn*, critical for corneodesmosome formation^36^. Increased expression of *H2-D1*, an HLA gene involved in antigen presentation^37^, suggests a role in early antigen response. Changes were also observed in *Alox12b* and *Serpinb5,* related to known EoE dysregulated genes^9, 38^. Interestingly, *Dsc1* and *Dsg3* showed increased expression, possibly responding to *Dsg1* loss^39^. The identified gene changes closely associated with the loss of *Ikk*β in EoE development closely mirror known alterations observed in the esophageal epithelium of EoE patients and moreover highlight new candidates for further investigation, establishing the *Ikk*β-deficient mouse model as a valuable tool for dissecting epithelial features and remodeling in the context of EoE. These gene changes mirror known alterations in the esophageal epithelium of EoE patients, highlighting new candidates for further investigation. This establishes the *Ikk*β-deficient mouse model as a valuable tool for dissecting epithelial features and remodeling in EoE.

## Discussion

EoE is a complex disorder characterized by allergic responses to food allergens, leading to chronic Th2-mediated inflammation, esophageal eosinophilia, and tissue remodeling and ultimately resulting in esophageal dysmotility, dysphagia, and food impaction. Due to the multifactorial nature of EoE, mouse models are indispensable for elucidating early disease events, the pathophysiological timeline, and therapeutic approaches^40^. After observing decreased IKKβ/NFκB signaling in esophageal mucosal cells in human EoE, we generated mice with a conditional knockout of esophageal epithelial *Ikk*β to investigate its role in pathogenesis.

Our analyses showed significant similarities between human EoE and the *Ikkβ^EEC-KO^/EoE* mouse model in both histological and transcriptional aspects. *Ikkβ^EEC-KO^/EoE* mice developed pronounced EoE features, such as eosinophil infiltration, intraepithelial eosinophils, microabscesses, DIS, BCH, CD4+ T cell recruitment, and lamina propria remodeling. This was accompanied by activation of IL-4/IL-13 signaling, innate immune signaling, and interferon signaling, pathways also observed in human EoE^41–43^. We also observed decreased expression of genes related to keratinocyte differentiation and skin barrier integrity, alongside increased proliferation. These findings align with those observed in human EoE^30, 44–46^.

Our mouse model demonstrates that altered epithelial signaling in response to an allergy trigger can replicate EoE features, confirming the esophageal epithelium as a key driver of EoE pathogenesis^9^. Given the central role of NF-κB in inflammatory responses, our findings are particularly intriguing. NF-κB has multifaceted, context- and cell-dependent functions, acting as a crucial signaling molecule in inflammation, epithelial proliferation, tissue dynamics, barrier integrity, and epithelial-immune homeostasis^47^. In the lower gut, NF-κB signaling in epithelial cells is essential for maintaining epithelial barrier integrity and microflora-epithelial homeostasis^48^. Specifically, inhibiting NF-κB signaling through conditional *Ikk*β knockout results in chronic inflammation, impaired barrier function, bacterial translocation into the mucosa, innate immune activation, and T-cell mediated inflammation^48^.

This study allowed us to identify genes potentially directly regulated by *Ikk*β loss relevant to EoE progression. Among those genes were *Dsg1a*, a desmosomal cadherin implicated as a loss-of-function risk factor for EoE development^49^, which is down-regulated downstream of IL-13 signaling in EoE^34^. This is accompanied by the loss of other key epithelial barrier proteins contributing to desmosome function, such as *Ppl*, encoding periplakin, which links desmosomes to intermediate filaments, and Cdsn, encoding corneodesmosin, a key structural component of corneodesmosomes in terminally differentiated tissue^36^. This indicates impaired desmosome formation and function in the esophageal epithelium of *Ikkβ^EEC-KO^*/EoE mice, suggesting a loss of cell-cell adhesion leading to an impaired barrier and potential consequences on the regulation of desmosome-directed differentiation.

In addition to genes contributing to the epithelial barrier, our cohort includes genes linked to epithelial *Ikk*β loss in EoE development, showing decreased expression of epithelial differentiation genes such as *Krt78*, involved in terminal differentiation and known to be lost in human EoE^35^. The cohort also includes the increased expression of *Serpinb5*, aligning with the dysregulation of serine protease signaling known to be a hallmark of EoE^9^. Additionally, we observed increased expression of the MHC class I component *H2-D1*, reflecting the increase in HLA gene expression in human EoE esophageal epithelium, supporting a role for the epithelium in early antigen presentation and immune response induction^50^. Moreover, the gene cohort includes up-regulation of *Hmgn1*, a high mobility group protein that regulates chromatin compaction^51^, histone modification^51^, and gene transcription^52^. *Hmgn1* has been implicated in epithelial differentiation, cellular adhesion, and p63 expression in other systems^53^.

Future studies will dissect early disease events combined with epithelial fate-mapping to understand disease progression and epithelial cell-autonomous and non-cell autonomous events in *Ikkβ^EEC-KO^/EoE* mice. This model will serve as a platform for testing epithelial-specific disease targets for EoE intervention. Lastly, although our study primarily focused on the esophageal epithelial transcriptome, future research will explore ligand-receptor crosstalk between epithelial subsets and other cell types, providing a comprehensive understanding of non-epithelial cell contributions to EoE progression.

### Conclusions

We have generated a novel mouse model with loss of epithelial IKKβ signaling in the presence of an allergic trigger. This model offers significant potential for developing strategies to prevent long-term chronic inflammation and tissue remodeling in EoE.

## Abbreviations

BCH: basal cell hyperplasia
DEGs: differentially expressed genes
DIS: dilation of intercellular spaces
EEC: esophageal epithelial cells
EoE: eosinophilic esophagitis
GSEA: gene set enrichment analysis
HC: healthy controls
IKBIP: IKKβ interacting protein gene
LogFC: log fold change
OVA: ovalbumin
scRNA-seq: single cell RNA sequencing

## References

1. Blanchard C, Mingler MK, Vicario M, Abonia JP, Wu YY, Lu TX, et al. IL-13 involvement in eosinophilic esophagitis: transcriptome analysis and reversibility with glucocorticoids. J Allergy Clin Immunol 2007; 120:1292–300.

2. Dellon ES, Woosley JT, Arrington A, McGee SJ, Covington J, Moist SE, et al. Rapid Recurrence of Eosinophilic Esophagitis Activity After Successful Treatment in the Observation Phase of a Randomized, Double-Blind, Double-Dummy Trial. Clin Gastroenterol Hepatol 2020; 18:1483–92 e2.

3. Greuter T, Bussmann C, Safroneeva E, Schoepfer AM, Biedermann L, Vavricka SR, et al. Long-Term Treatment of Eosinophilic Esophagitis With Swallowed Topical Corticosteroids: Development and Evaluation of a Therapeutic Concept. Am J Gastroenterol 2017; 112:1527–35.

4. Ingerski LM, Modi AC, Hood KK, Pai AL, Zeller M, Piazza-Waggoner C, et al. Health-related quality of life across pediatric chronic conditions. J Pediatr 2010; 156:639–44.

5. Safroneeva E, Coslovsky M, Kuehni CE, Zwahlen M, Haas NA, Panczak R, et al. Eosinophilic oesophagitis: relationship of quality of life with clinical, endoscopic and histological activity. Aliment Pharmacol Ther 2015; 42:1000–10.

6. Capocelli KE, Fernando SD, Menard-Katcher C, Furuta GT, Masterson JC, Wartchow EP. Ultrastructural features of eosinophilic oesophagitis: impact of treatment on desmosomes. J Clin Pathol 2015; 68:51–6.

7. Collins MH. Histopathologic features of eosinophilic esophagitis. Gastrointest Endosc Clin N Am 2008; 18:59–71; viii-ix.

8. Katzka DA, Ravi K, Geno DM, Smyrk TC, Iyer PG, Alexander JA, et al. Endoscopic Mucosal Impedance Measurements Correlate With Eosinophilia and Dilation of Intercellular Spaces in Patients With Eosinophilic Esophagitis. Clin Gastroenterol Hepatol 2015; 13:1242–8 e1.

9. Rochman M, Azouz NP, Rothenberg ME. Epithelial origin of eosinophilic esophagitis. J Allergy Clin Immunol 2018; 142:10–23.

10. Collins MH, Martin LJ, Alexander ES, Boyd JT, Sheridan R, He H, et al. Newly developed and validated eosinophilic esophagitis histology scoring system and evidence that it outperforms peak eosinophil count for disease diagnosis and monitoring. Dis Esophagus 2017; 30:1–8.

11. Laky K, Kinard JL, Li JM, Moore IN, Lack J, Fischer ER, et al. Epithelial-intrinsic defects in TGFbetaR signaling drive local allergic inflammation manifesting as eosinophilic esophagitis. Sci Immunol 2023; 8:eabp9940.

12. Bollrath J, Greten FR. IKK/NF-kappaB and STAT3 pathways: central signalling hubs in inflammation-mediated tumour promotion and metastasis. EMBO Rep 2009; 10:1314–9.

13. Grivennikov SI, Karin M. Dangerous liaisons: STAT3 and NF-kappaB collaboration and crosstalk in cancer. Cytokine Growth Factor Rev 2010; 21:11–9.

14. Oeckinghaus A, Ghosh S. The NF-kappaB family of transcription factors and its regulation. Cold Spring Harb Perspect Biol 2009; 1:a000034.

15. Pahl HL. Activators and target genes of Rel/NF-kappaB transcription factors. Oncogene 1999; 18:6853–66.

16. Chae S, Eckmann L, Miyamoto Y, Pothoulakis C, Karin M, Kagnoff MF. Epithelial cell I kappa B-kinase beta has an important protective role in Clostridium difficile toxin A-induced mucosal injury. J Immunol 2006; 177:1214–20.

17. Chen LW, Chen PH, Chang WJ, Wang JS, Karin M, Hsu CM. IKappaB-kinase/nuclear factor-kappaB signaling prevents thermal injury-induced gut damage by inhibiting c-Jun NH2-terminal kinase activation. Crit Care Med 2007; 35:1332–40.

18. Eckmann L, Nebelsiek T, Fingerle AA, Dann SM, Mages J, Lang R, et al. Opposing functions of IKKbeta during acute and chronic intestinal inflammation. Proc Natl Acad Sci U S A 2008; 105:15058–63.

19. Hsu LC, Enzler T, Seita J, Timmer AM, Lee CY, Lai TY, et al. IL-1beta-driven neutrophilia preserves antibacterial defense in the absence of the kinase IKKbeta. Nat Immunol; 12:144–50.

20. Pasparakis M, Courtois G, Hafner M, Schmidt-Supprian M, Nenci A, Toksoy A, et al. TNF-mediated inflammatory skin disease in mice with epidermis-specific deletion of IKK2. Nature 2002; 417:861–6.

21. Stratis A, Pasparakis M, Markur D, Knaup R, Pofahl R, Metzger D, et al. Localized inflammatory skin disease following inducible ablation of I kappa B kinase 2 in murine epidermis. J Invest Dermatol 2006; 126:614–20.

22. Li ZW, Omori SA, Labuda T, Karin M, Rickert RC. IKK beta is required for peripheral B cell survival and proliferation. J Immunol 2003; 170:4630–7.

23. Tetreault MP, Yang Y, Travis J, Yu QC, Klein-Szanto A, Tobias JW, et al. Esophageal squamous cell dysplasia and delayed differentiation with deletion of kruppel-like factor 4 in murine esophagus. Gastroenterology 2010; 139:171–81 e9.

24. Noti M, Wojno ED, Kim BS, Siracusa MC, Giacomin PR, Nair MG, et al. Thymic stromal lymphopoietin-elicited basophil responses promote eosinophilic esophagitis. Nat Med 2013; 19:1005–13.

25. Wiles KN, Alioto CM, Hodge NB, Clevenger MH, Tsikretsis LE, Lin FTJ, et al. IkappaB Kinase-beta Regulates Neutrophil Recruitment Through Activation of STAT3 Signaling in the Esophagus. Cell Mol Gastroenterol Hepatol 2021; 12:1743–59.

26. Patel CK, Kahrilas PJ, Hodge NB, Tsikretsis LE, Carlson DA, Pandolfino JE, et al. RNA-sequencing reveals molecular and regional differences in the esophageal mucosa of achalasia patients. Sci Rep 2022; 12:20616.

27. Greten FR, Eckmann L, Greten TF, Park JM, Li ZW, Egan LJ, et al. IKKbeta links inflammation and tumorigenesis in a mouse model of colitis-associated cancer. Cell 2004; 118:285–96.

28. Clevenger MH, Karami AL, Carlson DA, Kahrilas PJ, Gonsalves N, Pandolfino JE, et al. Suprabasal cells retain progenitor cell identity programs in eosinophilic esophagitis- driven basal cell hyperplasia. JCI Insight 2023; 8.

29. Kabir MF, Karami AL, Cruz-Acuna R, Klochkova A, Saxena R, Mu A, et al. Single cell transcriptomic analysis reveals cellular diversity of murine esophageal epithelium. Nat Commun 2022; 13:2167.

30. Rochman M, Wen T, Kotliar M, Dexheimer PJ, Ben-Baruch Morgenstern N, Caldwell JM, et al. Single-cell RNA-Seq of human esophageal epithelium in homeostasis and allergic inflammation. JCI Insight 2022; 7.

31. Tirosh I, Izar B, Prakadan SM, Wadsworth MH, 2nd, Treacy D, Trombetta JJ, et al. Dissecting the multicellular ecosystem of metastatic melanoma by single-cell RNA-seq. Science 2016; 352:189–96.

32. Kuleshov MV, Jones MR, Rouillard AD, Fernandez NF, Duan Q, Wang Z, et al. Enrichr: a comprehensive gene set enrichment analysis web server 2016 update. Nucleic Acids Res 2016; 44:W90–7.

33. Xie Z, Bailey A, Kuleshov MV, Clarke DJB, Evangelista JE, Jenkins SL, et al. Gene Set Knowledge Discovery with Enrichr. Curr Protoc 2021; 1:e90.

34. Sherrill JD, Kc K, Wu D, Djukic Z, Caldwell JM, Stucke EM, et al. Desmoglein-1 regulates esophageal epithelial barrier function and immune responses in eosinophilic esophagitis. Mucosal Immunol 2014; 7:718–29.

35. Kc K, Rothenberg ME, Sherrill JD. In vitro model for studying esophageal epithelial differentiation and allergic inflammatory responses identifies keratin involvement in eosinophilic esophagitis. PLoS One 2015; 10:e0127755.

36. Holthofer B, Windoffer R, Troyanovsky S, Leube RE. Structure and function of desmosomes. Int Rev Cytol 2007; 264:65–163.

37. Shiina T, Blancher A, Inoko H, Kulski JK. Comparative genomics of the human, macaque and mouse major histocompatibility complex. Immunology 2017; 150:127–38.

38. Matoso A, Allen D, Herzlinger M, Ferreira J, Chen S, Lu S, et al. Correlation of ALOX15 expression with eosinophilic or reflux esophagitis in a cohort of pediatric patients with esophageal eosinophilia. Hum Pathol 2014; 45:1205–12.

39. Godsel LM, Roth-Carter QR, Koetsier JL, Tsoi LC, Huffine AL, Broussard JA, et al. Translational implications of Th17-skewed inflammation due to genetic deficiency of a cadherin stress sensor. J Clin Invest 2022; 132.

40. Pilat JM, Jacobse J, Buendia MA, Choksi YA. Animal models of eosinophilic esophagitis. J Leukoc Biol 2024.

41. Avlas S, Shani G, Rhone N, Itan M, Dolitzky A, Hazut I, et al. Epithelial cell-expressed type II IL-4 receptor mediates eosinophilic esophagitis. Allergy 2023; 78:464–76.

42. Dellon ES, Rothenberg ME, Collins MH, Hirano I, Chehade M, Bredenoord AJ, et al. Dupilumab in Adults and Adolescents with Eosinophilic Esophagitis. N Engl J Med 2022; 387:2317–30.

43. Ruffner MA, Hu A, Dilollo J, Benocek K, Shows D, Gluck M, et al. Conserved IFN Signature between Adult and Pediatric Eosinophilic Esophagitis. J Immunol 2021; 206:1361–71.

44. Clevenger MH, Karami AL, Carlson DA, Kahrilas PJ, Gonsalves N, Pandolfino JE, et al. Suprabasal cells retain progenitor cell identity programs in eosinophilic esophagitis- driven basal cell hyperplasia. JCI Insight 2023.

45. Ding J, Garber JJ, Uchida A, Lefkovith A, Carter GT, Vimalathas P, et al. An esophagus cell atlas reveals dynamic rewiring during active eosinophilic esophagitis and remission. Nat Commun 2024; 15:3344.

46. Jiang M, Ku WY, Zhou Z, Dellon ES, Falk GW, Nakagawa H, et al. BMP-driven NRF2 activation in esophageal basal cell differentiation and eosinophilic esophagitis. J Clin Invest 2015; 125:1557–68.

47. Pasparakis M. Role of NF-kappaB in epithelial biology. Immunol Rev 2012; 246:346–58.

48. Nenci A, Becker C, Wullaert A, Gareus R, van Loo G, Danese S, et al. Epithelial NEMO links innate immunity to chronic intestinal inflammation. Nature 2007; 446:557–61.

49. Davis BP, Stucke EM, Khorki ME, Litosh VA, Rymer JK, Rochman M, et al. Eosinophilic esophagitis-linked calpain 14 is an IL-13-induced protease that mediates esophageal epithelial barrier impairment. JCI Insight 2016; 1:e86355.

50. Mulder DJ, Pooni A, Mak N, Hurlbut DJ, Basta S, Justinich CJ. Antigen presentation and MHC class II expression by human esophageal epithelial cells: role in eosinophilic esophagitis. Am J Pathol 2011; 178:744–53.

51. Lim JH, Catez F, Birger Y, West KL, Prymakowska-Bosak M, Postnikov YV, et al. Chromosomal protein HMGN1 modulates histone H3 phosphorylation. Mol Cell 2004; 15:573–84.

52. Bustin M. Chromatin unfolding and activation by HMGN(*) chromosomal proteins. Trends Biochem Sci 2001; 26:431–7.

53. Birger Y, Davis J, Furusawa T, Rand E, Piatigorsky J, Bustin M. A role for chromosomal protein HMGN1 in corneal maturation. Differentiation 2006; 74:19–29.

