## Supplemental figure legend for "Esophageal epithelial *Ikkβ* deletion promotes eosinophilic esophagitis in experimental allergy mouse model"

**Supplemental Figure 1.** **Sensitization, intraesophageal allergic challenge and esophageal epithelial *Ikkβ* loss are necessary for the experimental EoE protocol to recapitulate all key histological features of human disease.** (**A**) Hematoxylin and Eosin staining of esophageal sections of control or *Ikkβ^EEC-KO^* treated with the following conditions: no treatment, vehicle treatment, sensitization only treatment, or treatment with the EoE protocol. (**B**) Quantification of EoE-HSS disease activity scoring (**B**) for control or *Ikkβ^EEC-KO^* mice treated with the following treatment conditions: no treatment, vehicle treatment, sensitization only treatment, or treatment with the EoE protocol.

**Supplemental Figure 2. Lamina propria fibrosis is observed in experimental EoE with esophageal epithelial *Ikkβ* loss.** (**A-C**) Masson’s Trichrome staining of esophageal sections of *Ikkβ^EEC-KO^* (**A**) and *Ikkβ^EEC-KO^*/EoE mice (**B**). (**C**) Quantification of fibrosis scoring. Bar graphs represent means ± SEM. All statistics were determined by a 2-tailed Student t-test. Scale bar: 50 µm. n= 6 mice per experimental group. ***P* ≤ 0.01.

**Supplemental Figure 3. Contribution of sensitization, intraesophageal allergic challenge and esophageal epithelial *Ikkβ* loss to individual features of the EoE-HSS. (A-G**) Quantification of the individual features of EoE-HSS disease activity scoring for control or *Ikkβ^EEC-KO^* mice treated with the following treatment conditions: no treatment, vehicle treatment, sensitization only treatment, or treatment with the EoE protocol. Individual features include total eosinophil influx score (**A**), intraepithelial influx score (**B**), eosinophil abscess score (**C**), BCH score (**D**), dyskeratotic epithelial cell score (**E**), dilated intercellular spaces score (**F**) and fibrosis score (**G**). All statistics were determined by a two-way ANOVA with post-hoc Tukey’s multiple comparisons test and calculation of partial omega squared. Scale bar: 50 µm. n= 6 mice per experimental group. **P* ≤ 0.05, ***P* ≤ 0.01, ****P* ≤ 0.001, *****P* ≤ 0.0001.

**Supplemental Figure 4. Characteristics of esophageal mucosal cells in *Ikkβ^EEC-KO^* and *Ikkβ^EEC-KO^*/EoE mice.** (**A**) Heatmap of the top genes expressed per cell type. Ranked by average logFC, FDR < 0.05, row-normalized expression z-scores**.** (**B**) UMAP showing esophageal mucosal cells from *IKKβ^EEC-KO^* and *IKKβ^EEC-KO^*/EoE mice. (**C**) Bar plot showing the frequency of *Ikkβ^EEC-KO^* and *Ikkβ^EEC-KO^*/EoE biological replicates per each esophageal mucosal cell type.

**Supplemental Figure 5. Clustering and annotation of the epithelial murine EoE dataset.** (**A**) UMAP of the unbiased clustering of EEC from the murine EoE dataset. (**B**) Violin plots displaying the expression of established gene markers for dividing, quiescent, basal, and differentiating EEC. (**C**) Clustering tree analysis showing the nearest neighbor clustering on the epithelial dataset using 30 principal compartments across resolutions 0-0.18. Basal clusters were isolated (dash circled and arrow) and re-clustered to obtain higher resolution. Dot size reflects the cell count per cluster, arrow transparency reflects the proportion of cells moving to the indicated cluster(s) in the next resolution, dot color reflects the cell compartment identity, star represents the selected resolution for clustering. (**D**) UMAP of the re-clustered basal epithelial cell dataset colored by identified basal clusters. (**E**) Violin pots showing the expression of gene markers of quiescence, S-phase, G2/M phase, and general basal cell identity within the re-clustered basal cell dataset. (**F**) Bar plot showing the frequency of *Ikkβ^EEC-KO^* and *Ikkβ^EEC-KO^*/EoE biological replicates per each EEC cluster. (**G**) Bar plot showing the frequency of each mouse group according to genotype and treatment condition per EEC cluster.

**Supplemental Figure 6. Gene expression profile across EEC clusters in the murine EoE dataset.** Heatmap of the top 5 genes expressed per cell type in the murine EoE dataset. Ranked by average logFC, FDR < 0.05, row-normalized expression z-scores**.**

**Supplemental Figure 7. Increased CD4+ T cells are detected in *Ikkβ^EEC-KO^*/EoE mice** (**A**) Violin plot showing the expression of *Cd4* in T cells across the different mouse groups within the murine EoE dataset. (**B**) Immunohistochemical staining against CD4 in esophageal sections of *Ikkβ^EEC-KO^* and *Ikkβ^EEC-KO^*/EoE mice.

**Supplemental Figure 8. Permutation test results. (A)** Range plot outcome for permutation tests conducted in Figure 4C. Horizontal bars represent the range of observed log2FC (obs_log2FD) values lie within 97.5% confidence intervals for the comparison of each cluster or compartment between *Ikkβ^EEC-KO^*/EoE and *Ikkβ^EEC-KO^* mice, with the dot indicating the observed Log2FC. Significance was determined by a false discovery rate (FDR) < 0.05 and |log2FC| difference > 0.58. (**B**) Range plot outcome for permutation tests conducted in Figure 4E. Horizontal bars represent the range of observed log2FC (obs_log2FD) values lie within 97.5% confidence intervals for the comparison of cell cycle stage proportions between *Ikkβ^EEC-KO^*/EoE and *Ikkβ^EEC-KO^* mice, with the dot indicating the mean observed Log2FC Significance was determined by a false discovery rate (FDR) < 0.05 and |log2FC| difference > 0.58.

**Supplemental Figure 9. Immunohistochemical staining against Ki67 for esophageal sections of *Ikkβ^EEC-KO^* and *Ikkβ^EEC-KO^*/EoE mice.** Scale bar: 50 µm. n= 6

**Supplemental Figure 10. Ridge plot showing the pseudotime value distribution across all EEC for each biological replicate of *Ikkβ^EEC-KO^* or *Ikkβ^EEC-KO^*/EoE mice.**

**Supplemental Figure 11. Expression profile of EoE-related genes closely associated with the loss of esophageal epithelial *Ikkβ*.** Heatmap of log_2_ normalized z-score expression of the 62 overlapping DEGs identified in Figure 6C across EEC compartments in *Ikkβ^EEC-KO^*, control/EoE or *Ikkβ^EEC-KO^*/EoE mice. Top hierarchical clusters are displayed.
