## Supplemental materials and methods for "Esophageal epithelial *Ikkβ* deletion promotes eosinophilic esophagitis in experimental allergy mouse model"

***Generation of ED-L2-Cre; Ikkβ^L/L^ Mice.***

All animal studies were approved by the Institutional Animal Care and Use Committee (IACUC) at Northwestern University. To generate *ED-L2-Cre; Ikkβ^L/L^* mice*, Ikkβ^L/L^* mice containing a floxed third exon of *Ikkβ*^1^ were crossed with mice harboring *Cre* recombinase under the control of the *EBV-ED-L2* promoter^2^. All mice used for experiments were on a pure Balb/C background.  For all experiments with *ED-L2-Cre; Ikkβ^L/L^* mice (ref: *Ikkβ^EEC-KO^*), sex-matched littermate *Ikkβ^L/L^* mice (ref: control) lacking the Cre transgene served as controls. Both male and female experimental groups were included. Mice were injected with BrdU 1h prior to sacrifice. Mice were also examined grossly for the presence of food impaction and processed for histology. Briefly, tissue was fixed in neutral buffered formalin (Thermo Fisher Scientific) for 24 hours and embedded in paraffin, and 4-μm sections were applied to charged plus slides. Slides were stained with hematoxylin and eosin, and images were captured on a Nikon Eclipse Ci-E microscope with a Nikon DS-Ri2 camera and NIS Elements software. The following numbers of age-matched and sex-matched mice were examined histologically: No treatment: 6 controls, 6 *Ikkβ^EEC-KO^* mice; vehicle treatment: 6 controls, 6 *Ikkβ^EEC-KO^* mice; ear sensitization only: 6 controls, 6 *Ikkβ^EEC-KO^* mice; MC903+OVA/OVA: 6 controls, 6 *Ikkβ^EEC-KO^* mice.

***Induction of EoE-like disease in mice.***

EoE-like disease was induced in mice using a methodology previously described^3^. Three to four month-old *Ikkβ^EEC-KO^* mice and littermate controls were treated topically on the ears for 12 days with 2nmol of MC903 (Tocris Bioscience) and 100 μg of OVA (Sigma Aldrich, St-Louis, MO), once daily. On day 15 and 17, mice were challenged intraesophageally (IE) with 50 mg OVA. Following the first IE OVA gavage, mice were given access to water containing 1.5g/L OVA ad libitum until sacrifice at day 18. For vehicle treatment, *Ikkβ^EEC-KO^* mice and littermate genetic controls were treated on the ears with a combination of ethanol and PBS and were gavaged with water. For sensitization only treatment, *Ikkβ^EEC-KO^* mice and littermate genetic controls were treated on the ears with a combination of MC903 and OVA and gavaged with water. Mice were weighed weekly and body condition was scored and recorded.

***Immunohistochemistry, Immunofluorescence and scoring.***

For formalin-fixed paraffin-embedded esophageal sections, heat antigen retrieval (2100 Antigen Retriever, Electron Microscopy Sciences, Hatfield, PA) was performed as previous described^4^. Primary and secondary antibodies were added, and detection was performed as previously described ^4^. For immunohistochemistry, biotinylated conjugated secondary antibodies were used. For fluorescent labeling, Alexa Fluor^TM^ 488 and Alexa Fluor^TM^ 568 (Thermo Fisher Scientific) conjugated secondary antibodies were used. Dapi was used as a nuclear stain. A detailed list of the antibodies used for staining is shown in **Supplemental Table 2**. Images were acquired on a Nikon Eclipse Ci microscope with a Nikon DS-Ri2 camera and NIS Elements software. H&E staining was performed by the Robert H. Lurie Comprehensive Cancer Center Pathology Core. Image analysis was performed using Fiji software (74). H&E-stained slides were evaluated for disease activity according to EoE-HSS, which is comprised of eight parameters: eosinophil infiltration (EI), eosinophil abscesses (EA), eosinophil surface layering (SL), dilated intercellular spaces (DIS), basal zone hyperplasia (BZH), dyskeratotic epithelial cells (DEC), and fibrosis (F) (11). The EoEHSS was adapted for mouse histology by exclusion of the surface epithelial alteration and by adapting fibrosis scoring.

***Western Blots.***

Cells were harvested in Triton lysis Buffer (1% Triton X-100, 50mM Tris-HCl pH 7.5, 100mM NaCl, 5mM EDTA, 40mM β-glycerophosphate, 5% glycerol, 50mM NaF) plus protease (Pierce, Rockford, IL) and phosphatase inhibitors (Sigma-Aldrich) and protein concentration was determined using Pierce BCA protein assay (Thermo Fisher Scientific). Proteins were separated on NuPage 4–12% Bis-Tris gels (Thermo Fisher Scientific) and transferred onto polyvinylidene difluoride membrane (EMD Millipore, Billerica, MA) or Nitrocellulose membranes (Biorad, Hercules, CA). After blocking, membranes were incubated overnight at 4°C with the following antibodies 1:800 rabbit phospho-p65 NFκB (Ser 536, #3033, Cell Signaling), 1:6,000 rabbit p65 NFκB (#8242, Cell Signaling), 1:10,000 rabbit anti-GAPDH (#G9545, Sigma-Aldrich). Quantitation of bands was performed with FIJI (version 2.0.0). Mean gray value was measured for each protein of interest and the values for phosphorylated proteins were normalized to total protein values.

***Human specimen sample collection and processing.***

Esophageal mucosal biopsies were collected from EoE patients and healthy controls (HC) during sedated endoscopy through the Digestive Health Foundation Biorepository. HC were healthy, asymptomatic, adult volunteers and met asymptomatic criteria including the lack of esophageal symptoms (heartburn, dysphagia, chest pain), history of tobacco use or alcohol dependency, BMI greater than 30 kg/m2, or previous treatment with antacids or proton pump inhibitors. 8 HC were enrolled and 19 EoE patients were recruited at the primary visit contingent upon confirmed diagnosis and no history of steroid treatment. Exclusion criteria for EoE included active severe esophagitis (Los Angeles esophagitis Grade C and above)^5^, evidence of mechanical obstruction due to peptic stricture (GERD), long-segment Barrett’s metaplasia, unstable medical illness with ongoing diagnostic workup and treatment, current drug or alcohol abuse or dependency, current neurologic or cognitive impairment that would make the patient an unsuitable candidate for a research trial, severe mental illness, pregnancy and bleeding diathesis, or need for anticoagulation that cannot be stopped for endoscopy. Biospecimen Reporting for improved study quality data including age, sex, and race is detailed in **Supplemental Table 3**. All procedures using human tissue received approval from the Northwestern Institutional Review Board (STU00208111) and all methods were performed in accordance with the relevant guidelines and regulations. Informed consent was obtained from all subjects/legal guardians prior to participation. Biopsies were collected from the proximal and distal esophagus, at 5 and 15 cm above the squamocolumnar junction, respectively. Biopsies were collected in RNA later (Thermo Fisher Scientific, Pittsburgh, PA) and stored at -80 °C until RNA extraction.

***Bulk RNA sequencing.***

Following homogenization in RLT buffer, RNA was extracted and purified using the RNeasy kit (Qiagen, Germantown, MD). Reverse transcription was performed with the Maxima First-Strand cDNA Synthesis for RT-qPCR kit (Thermo Fisher Scientific). For bulk RNA-sequencing, DNA libraries were generated using the Tru-Seq Stranded mRNA-seq library prep (Illumina, San Diego, CA) following the manufacturer’s instructions. Sequencing was performed using Illumina HiSeq 4000 (Northwestern University NuSeq Core Facility). The quality of DNA reads, in fastq format, was evaluated using FastQC. Adapters were trimmed and reads of poor quality or aligning to rRNA sequences were filtered. The cleaned reads were aligned to the Homo sapiens genome (hg38) using STAR38^6^. In conjunction with a gene annotation file for hg38 obtained from UCSC (University of California Santa Cruz; [http://genome.ucsc.edu](http://genome.ucsc.edu/)), read counts for each gene were calculated using HTSeq-count[^39^](https://www.ncbi.nlm.nih.gov/pmc/articles/PMC9712691/#CR39). DESeq2 was used to determine normalization and differential expression[^40^](https://www.ncbi.nlm.nih.gov/pmc/articles/PMC9712691/#CR40). The cutoff for determining significantly differentially expressed genes was an FDR-adjusted p-value less than 0.05. RNA-seq was deposited in Gene Expression Omnibus (GEO #270219, 271128) and can be accessed at <http://www.ncbi.nlm.nih.gov/geo/query/acc.cgi?acc=GSE270219>, <http://www.ncbi.nlm.nih.gov/geo/query/acc.cgi?acc=GSE271128>. A multivariate principal component analysis (PCA) was carried out on the entire dataset to reduce data dimensionality and to assess clustering. Enriched gene ontology (GO) terms were identified using Metascape online platform^5^ and Gene Set Enrichment Analysis (GSEA) was conducted using clusterProfiler^6,7^ package in R. Differentially expressed gene lists were ranked by log-fold change and analyzed against the Biological Process database via org.Hs.eg.db^8^ R package. Gene set size restricted between 3 and 800, and 10,000 permutations used to estimate the enrichment score for each gene set.

***scRNA-sequencing.***

*scRNA-seq sample preparation, library preparation and sequencing.*

Mouse esophagi were harvested digested in digested in Dispase (Corning, Corning, NY) diluted in HBSS containing 10 uM HEPES 10 uM and 10 ug/mL DNase I at 37 ⁰C for 15 min with 1500 rpm agitation followed by mechanical separation of the epithelium from whole esophagus. Tissue was then minced and digested in 0.25% trypsin containing 10 uM HEPES and 10 ug/ml DNase I for 30 min at 37 ⁰C with agitation. The cell suspension was filtered through a 40-um strainer followed by 12 min and 6 min centrifugation at 500g at 4⁰C. Resuspended pellets were filtered through a 40 um flowmi filter (SP Bel-Art, Wayne, NJ), and measured for cell count and viability using the Cellometer Auto2000 (Nexcelom Bioscience, Lawrence, MA). All cell suspensions met an 85% minimum viability. Samples were sequenced as previously described in section 2.2.2.1, with the GRCm38 transcriptome used as reference for alignment and feature counting using Cell Ranger (V4.0.0/6.0.0/6.1.0, 10X Genomics).

*Data filtering, integration, and clustering.*

Filtered matrix files were processed as Seurat objects in the Seurat R package 4.2.0^9^ with a minimum threshold of expression in ≥5 cells per gene. Datasets were individually normalized, scaled, and processed to calculate variable features using Seurat’s SCTransform workflow. Individual filtered samples were then integrated using reverse principal component analysis dimensional reduction with a representative *IKKβ^ECC-KO^* sample used as reference. The Integrated dataset was filtered to exclude non-epithelial cells with unique gene counts <400, epithelial cells with unique gene counts <1000, and all cells with percent mitochondrial transcripts >20%. Dimensionality reduction was performed followed by calculation of UMAP embeddings, nearest neighbors, and graph-based clustering. Clusters were annotated according to the expression of known cell-specific gene markers and confirmed against the transcriptional profiles identified by Seurat’s function FindAllMarkers.

*Epithelial cluster and compartment identification.*

Epithelial cells were subsetted and reintegrated on a per-sample basis using the Seurat integration pipeline described above with integration anchors were calculated against a representative *Ikkβ^ECC-KO^* sample as reference. Principal component analysis (PCA) was performed and the first 30 PCs were included for downstream analysis. Optimal clustering resolution of 0.13 was determined using Clustree. Quiescent (clusters 1-2) and cycling clusters (clusters 3-5) were subclustered to distinguish quiescent basal cells from different cycling populations (S phase, G2/M phase, post-G2/M phase). Epithelial clusters were annotated according to expression of known genes in HC as previously described (25) and confirmed against the transcriptional profiles identified by FindAllMarkers, performed on HC cells. Clusters were combined into parental epithelial compartments (Basal, Suprabasal, Superficial) based on the expression of established markers^10–13^.

*Cell cycle and proliferation analysis.*

Seurat’s function CellCycleScoring^14^ was used to assign the cell cycle phase of each cell. Cells exhibiting a weak predicted score for S and G2/M were classified as G0/G1 phase. Expression of the markers *Krt15* and *Dst* identified Basal_1 and Basal_2 epithelial clusters as quiescent and distinguished G0 and G1 phase. During the SCTransform workflow, cell cycle was not regressed, allowing EEC to cluster based on quiescence, S-phase, G2/M-phase and progressive stages of differentiation, confirmed using the expression of marker genes and cell cycle scoring for each cluster. Cell proportion in each cluster was used to assess proliferation rates.

*Detection of DEGs, gene expression analysis and gene set enrichment analysis.*

Identification of DEGs between cell clusters was performed using FindAllMarkers, with filtering for significantly upregulated genes with logFC > 0.25. For differential expression analysis comparing expression profiles between like cell identities across disease conditions, Seurat’s FindMarkers function was employed using the MAST test^7^. DEGs were filtered based on an FDR-adjusted P value < 0.05 and |log2FC| > 0.25, unless a more stringent thresholding was specified. Pathway enrichment analyses were performed on DEGs filtered for logFC and significance based on FDR adjusted P value using Metascape^5^. Enrichment analysis for upstream transcription factor regulation was conducted with EnrichR^15,16^ using the Chea_2022 database^8^.

*Heatmap visualization, population z-score calculation and hierarchical clustering.*

Gene sets displayed in heatmaps were confirmed as changed in experimental groups with differential expression testing filtered based on FDR-adjusted P value < 0.05 and minimum logFC threshold. To calculate population z-scores, average population expression values were derived from the normalized RNA assay and scaled by the mean and standard deviation calculated across all populations. DEGs were clustered using k-means clustering and heatmaps were generated using the R package Complex Heatmap 2.10.0^17,18^.

*Gene signature score analysis and functional analysis.*

Gene signatures were generated using Seurat’s function AddModuleScore. Quiescent and superficial gene signatures were defined using cells from *Ikkβ^EEC-KO^* mice from our scRNA-seq dataset. Differential expression analysis was performed comparing either quiescent epithelial clusters (B1-2) or superficial clusters (SF1-SF2) as compared to the remaining epithelium. DEGs were filtered for FDR-adjusted P value < 0.05 and ranked by logFC, with the top 100 ranked genes selected.

*Pseudotime analysis.*

Pseudotime analysis was performed using the R package Monocle3 1.0.0^19^. Samples were log2 normalized, scaled, merged using Seurat’s merge function, dimensionally reduced, batch corrected using the FMNN method by individual sample, and UMAP embeddings were calculated. A CellDataSet object was created with normalized and scaled counts for 2000 variable genes and reduction feature loadings calculated by FMNN. Monocle3's function learn_graph was used to infer a trajectory graph from the UMAP embeddings, with Euclidean distance ratio of 1, geodesic distance ratio of 0.33, and a minimum branch length of 10. Cells within the S-phase epithelial cluster were assigned a root state of pseudotime 0. Increasing pseudotime values of cells committed to becoming quiescent are depicted to the left, and pseudotime values of cells committed to differentiation to the right on pseudotime axes.

*Data and code availability.*

All raw sequencing files and processed barcode and feature matrices used within the article are deposited in NCBI’s Gene Expression Omnibus (GEO) database under accession codes GSE270219 and GSE271128. All supporting analytic code is available at the 'scRNA-Mouse_EoE_Esophagus' repository hosted by the Tetreault Lab on GitHub.

***Statistical analyses.***

Statistical analyses were performed using R version 4.1.1. Descriptive statistics are displayed as mean ± standard error of the mean for continuous variables unless otherwise described and frequency counts for categorical variables. For the comparison of multiple conditions including genotype and treatment models, a two-way analysis of variance (ANOVA) test was employed. Post-hoc Tukey’s multiple comparisons test was subsequently applied to compare groups of means. Additionally, partial omega squared was calculated to quantify the proportion of variance in the dependent variable exclusively explained by each independent variable^20^. For the comparison of two normally distributed groups, a one-tailed Student's T-test was utilized. To compare proportions within the scRNA-seq dataset, we utilized the R package scProportionTest^21^. Permutation testing with 10,000 permutations was conducted to determine statistical significance, while bootstrapping was employed to calculate confidence intervals for observed changes. Statistical significance was defined as *P*-values < 0.05 and |logFC| > 0.58.

1.	Li ZW, Omori SA, Labuda T, Karin M, Rickert RC. IKK beta is required for peripheral B cell survival and proliferation. J Immunol 2003; 170:4630-7.

2.	Tetreault MP, Yang Y, Travis J, Yu QC, Klein-Szanto A, Tobias JW, et al. Esophageal squamous cell dysplasia and delayed differentiation with deletion of kruppel-like factor 4 in murine esophagus. Gastroenterology 2010; 139:171-81 e9.

3.	Noti M, Wojno ED, Kim BS, Siracusa MC, Giacomin PR, Nair MG, et al. Thymic stromal lymphopoietin-elicited basophil responses promote eosinophilic esophagitis. Nat Med 2013; 19:1005-13.

4.	Yang Y, Goldstein BG, Nakagawa H, Katz JP. Kruppel-like factor 5 activates MEK/ERK signaling via EGFR in primary squamous epithelial cells. FASEB J 2007; 21:543-50.

5.	Lundell LR, Dent J, Bennett JR, Blum AL, Armstrong D, Galmiche JP, et al. Endoscopic assessment of oesophagitis: clinical and functional correlates and further validation of the Los Angeles classification. Gut 1999; 45:172-80.

6.	Dobin A, Davis CA, Schlesinger F, Drenkow J, Zaleski C, Jha S, et al. STAR: ultrafast universal RNA-seq aligner. Bioinformatics 2013; 29:15-21.

7.	Finak G, McDavid A, Yajima M, Deng J, Gersuk V, Shalek AK, et al. MAST: a flexible statistical framework for assessing transcriptional changes and characterizing heterogeneity in single-cell RNA sequencing data. Genome Biol 2015; 16:278.

8.	Keenan AB, Torre D, Lachmann A, Leong AK, Wojciechowicz ML, Utti V, et al. ChEA3: transcription factor enrichment analysis by orthogonal omics integration. Nucleic Acids Res 2019; 47:W212-W24.
