## Supplemental figures for "Esophageal epithelial *Ikkβ* deletion promotes eosinophilic esophagitis in experimental allergy mouse model"

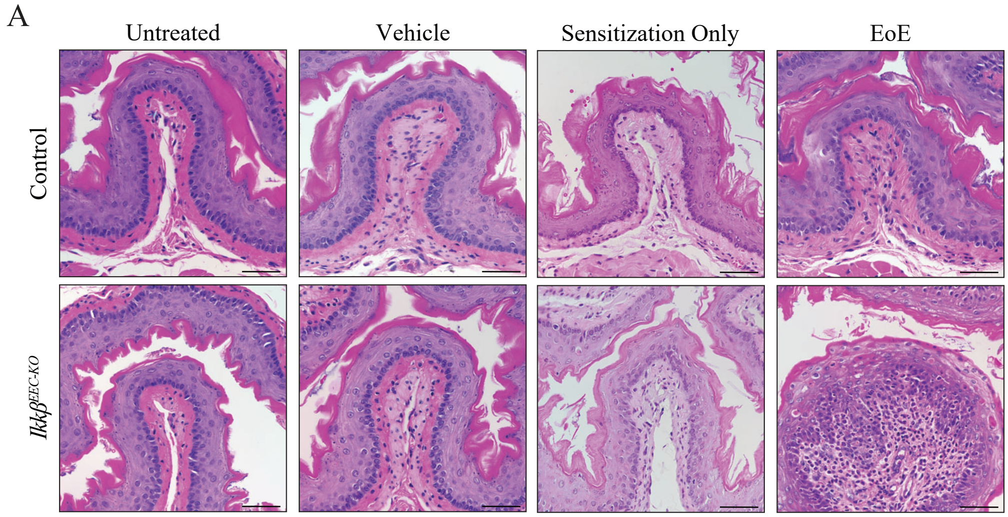


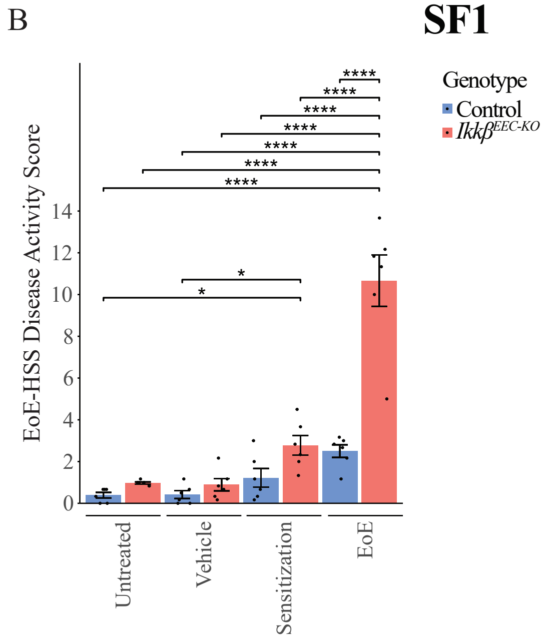


| | F-statistic | P value | $\omega^2_P$ |
|---|---|---|---|
| Genotype | 53.43 | 6.98e-09 | 0.15 |
| Treatment | 58.01 | 1.26e-14 | 0.50 |
| Genotype:Treatment | 24.92 | 2.94e-09 | 0.21 |

# SF2

A

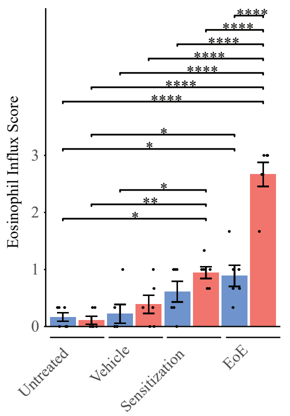


| | F-statistic | P value | $\omega^2_P$ |
|---|---|---|---|
| Genotype | 26.68 | 6.99e-06 | 0.10 |
| Treatment | 46.90 | 3.63e-13 | 0.54 |
| Genotype:Treatment | 14.89 | 1.17e-06 | 0.16 |

B

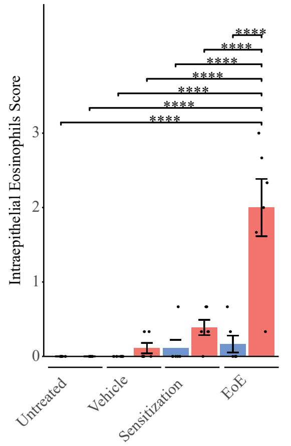


| | F-statistic | P value | $\omega^2_P$ |
|---|---|---|---|
| Genotype | 26.13 | 8.29e-06 | 0.14 |
| Treatment | 21.37 | 2.01e-8 | 0.34 |
| Genotype:Treatment | 15.64 | 7.00e-07 | 0.25 |

C

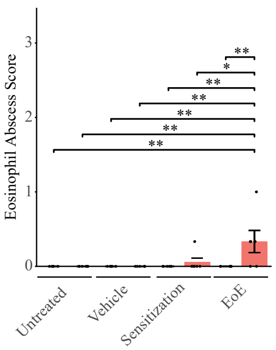


| | F-statistic | P value | $\omega^2_P$ |
|---|---|---|---|
| Genotype | 26.68 | 6.99e-06 | 0.10 |
| Treatment | 46.90 | 3.63e-13 | 0.54 |
| Genotype:Treatment | 14.89 | 1.17e-06 | 0.16 |

D

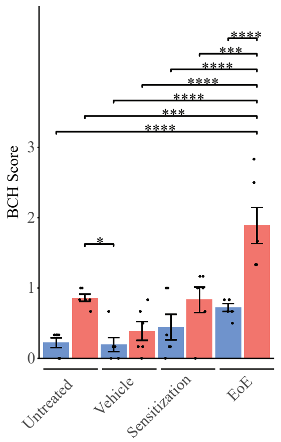


| | F-statistic | P value | $\omega^2_P$ |
|---|---|---|---|
| Genotype | 26.68 | 6.99e-06 | 0.10 |
| Treatment | 46.90 | 3.63e-13 | 0.54 |
| Genotype:Treatment | 14.89 | 1.17e-06 | 0.16 |

E

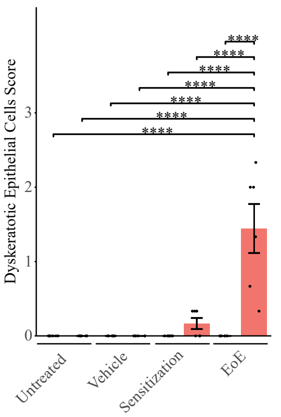


| | F-statistic | P value | $\omega^2_P$ |
|---|---|---|---|
| Genotype | 22.73 | 2.48e-05 | 0.13 |
| Treatment | 17.11 | 2.65e-07 | 0.29 |
| Genotype:Treatment | 17.11 | 2.65e-07 | 0.29 |

F

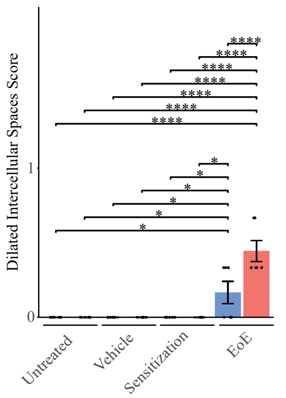


| | F-statistic | P value | $\omega^2_P$ |
|---|---|---|---|
| Genotype | 7.35 | 9.83e-03 | 0.034 |
| Treatment | 35.53 | 2.32e-11 | 0.59 |
| Genotype:Treatment | 7.35 | 4.90e-04 | 0.11 |

G

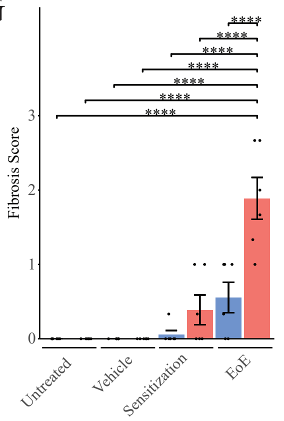


| | F-statistic | P value | $\omega^2_P$ |
|---|---|---|---|
| Genotype | 16.92 | 1.89e-04 | 0.09 |
| Treatment | 33.18 | 6.16e-11 | 0.52 |
| Genotype:Treatment | 9.70 | 6.15e-05 | 0.14 |

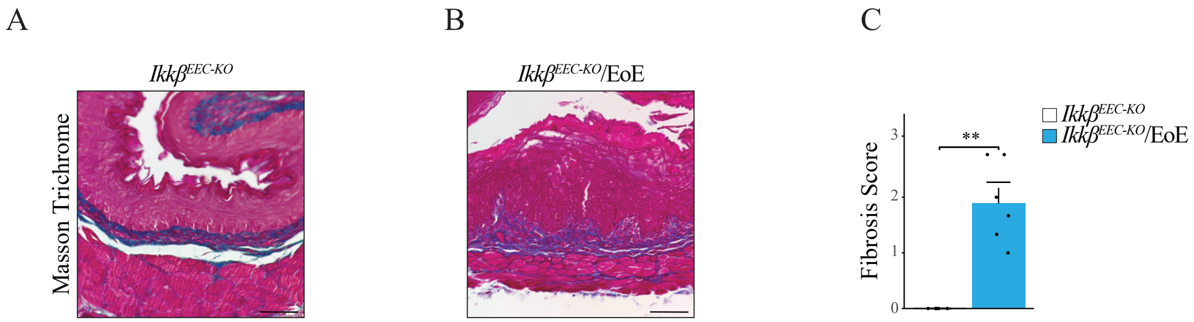
A
B
C
Ikkβ EEC-KO
Ikkβ EEC-KO/EoE
Masson Trichrome
Fibrosis Score
**
0
1
2
3
Ikkβ EEC-KO
Ikkβ EEC-KO/EoE

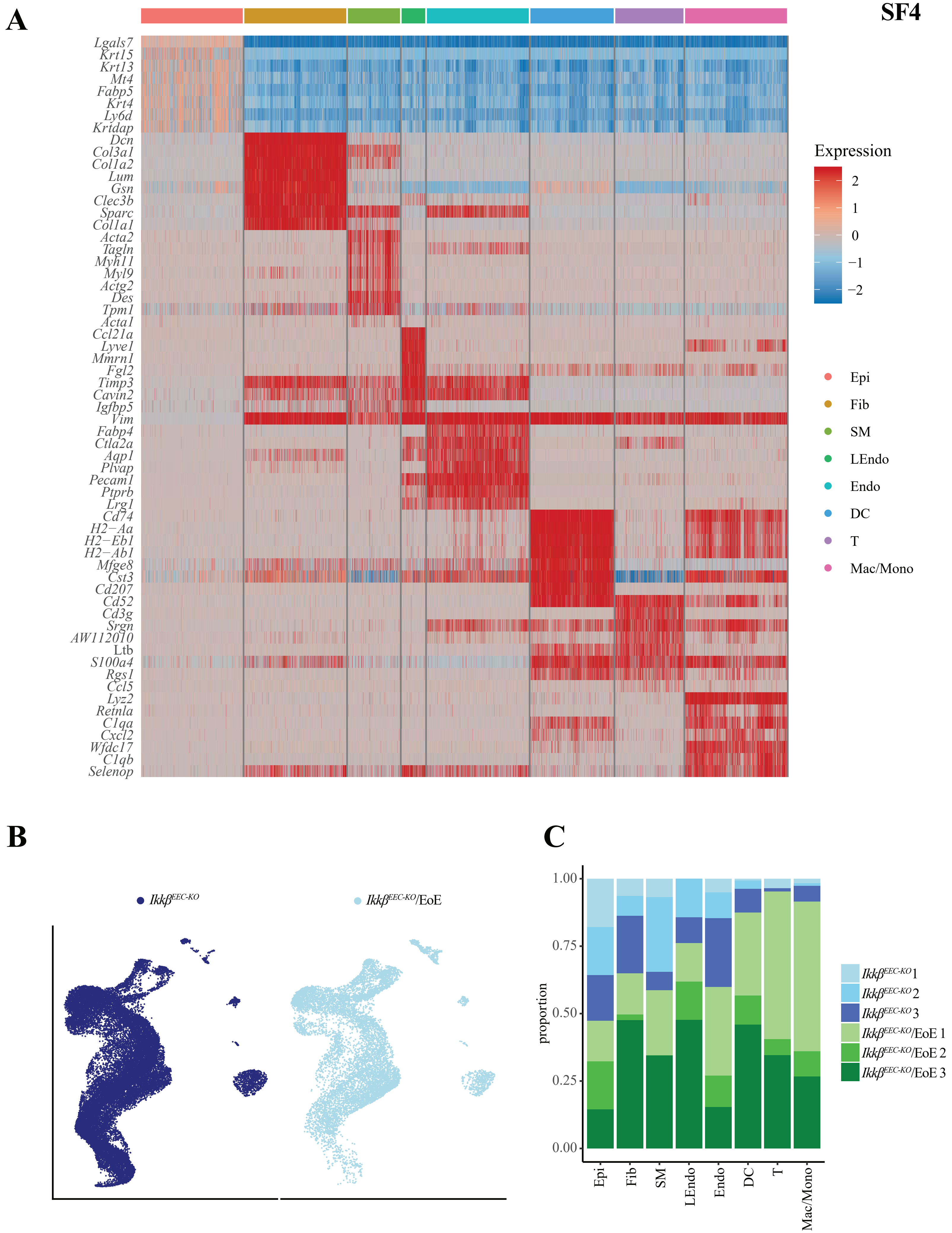
SF4
A
Lgals7
Krt15
Krt13
Mt4
Fabp5
Krt4
Ly6d
Krtdap
Dcn
Col3a1
Col1a2
Lum
Gsn
Clec3b
Sparc
Col1a1
Acta2
Tagln
Myh11
Myl9
Actg2
Des
Tpm1
Acta1
Ccl21a
Lyve1
Mmrn1
Fgl2
Timp3
Cavin2
Igfbp5
Vim
Fabp4
Ctla2a
Aqp1
Plvap
Pecam1
Ptprb
Lrg1
Cd74
H2−Aa
H2−Eb1
H2−Ab1
Mfge8
Cst3
Cd207
Cd52
Cd3g
Srgn
AW112010
Ltb
S100a4
Rgs1
Ccl5
Lyz2
Retnla
C1qa
Cxcl2
Wfdc17
C1qb
Selenop
Expression
2
1
0
−1
−2
Epi
Fib
SM
LEndo
Endo
DC
T
Mac/Mono
B
Ikkβ^EEC-KO
Ikkβ^EEC-KO/EoE
C
1.00
0.75
0.50
0.25
0.00
proportion
Epi
Fib
SM
LEndo
Endo
DC
T
Mac/Mono
Ikkβ^EEC-KO 1
Ikkβ^EEC-KO 2
Ikkβ^EEC-KO 3
Ikkβ^EEC-KO/EoE 1
Ikkβ^EEC-KO/EoE 2
Ikkβ^EEC-KO/EoE 3

**SF5**

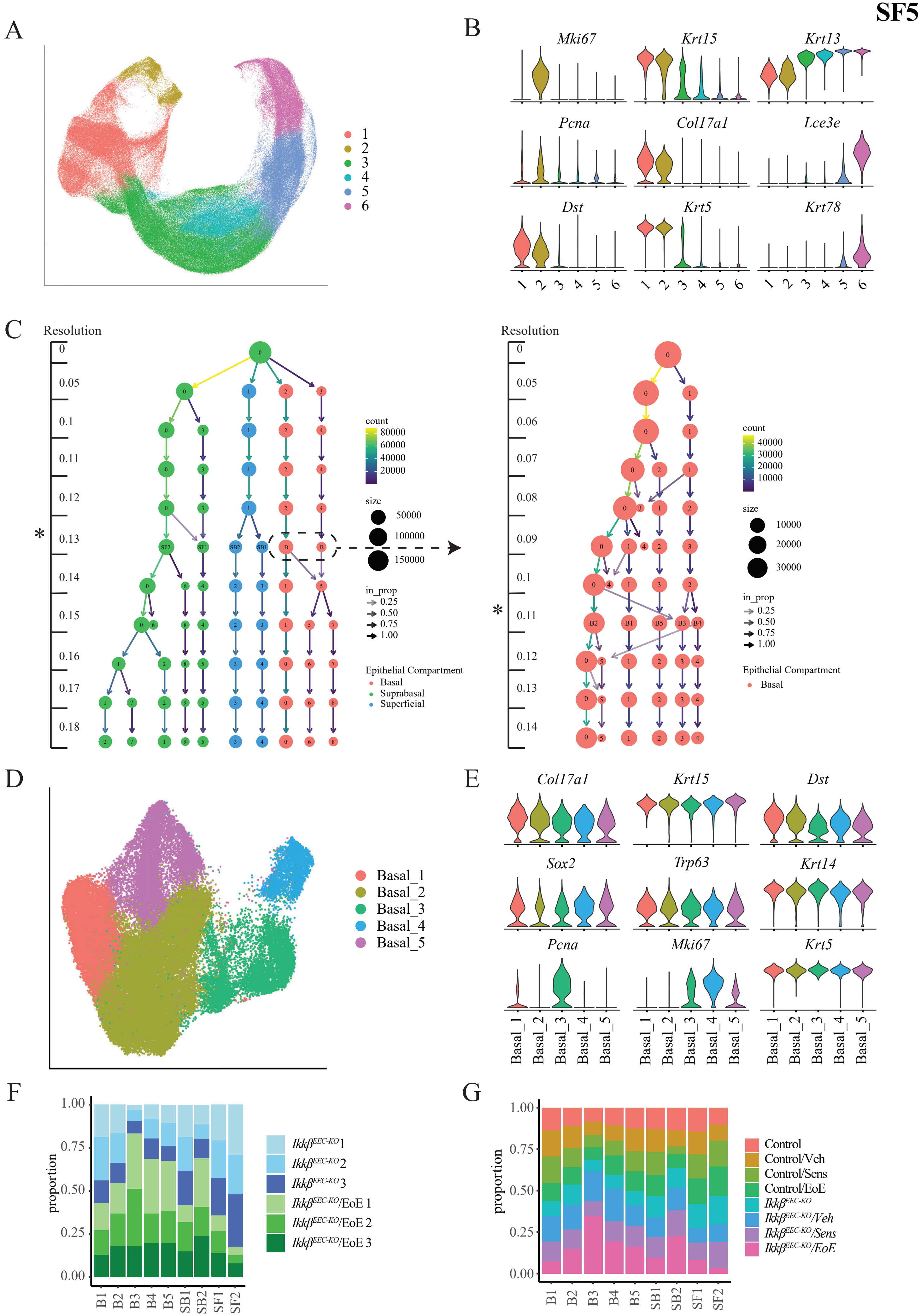
A
B
Mki67
Krt15
Krt13
Pcna
Col17a1
Lce3e
Dst
Krt5
Krt78
C
Resolution
count
size
in_prop
Epithelial Compartment
Basal
Suprabasal
Superficial
D
Basal_1
Basal_2
Basal_3
Basal_4
Basal_5
E
Sox2
Trp63
Krt14
F
proportion
Ikkβ^EEC-KO 1
Ikkβ^EEC-KO 2
Ikkβ^EEC-KO 3
Ikkβ^EEC-KO/EoE 1
Ikkβ^EEC-KO/EoE 2
Ikkβ^EEC-KO/EoE 3
G
Control
Control/Veh
Control/Sens
Control/EoE
Ikkβ^EEC-KO
Ikkβ^EEC-KO/Veh
Ikkβ^EEC-KO/Sens
Ikkβ^EEC-KO/EoE
B1
B2
B3
B4
B5
SB1
SB2
SF1
SF2

**SF6**

A

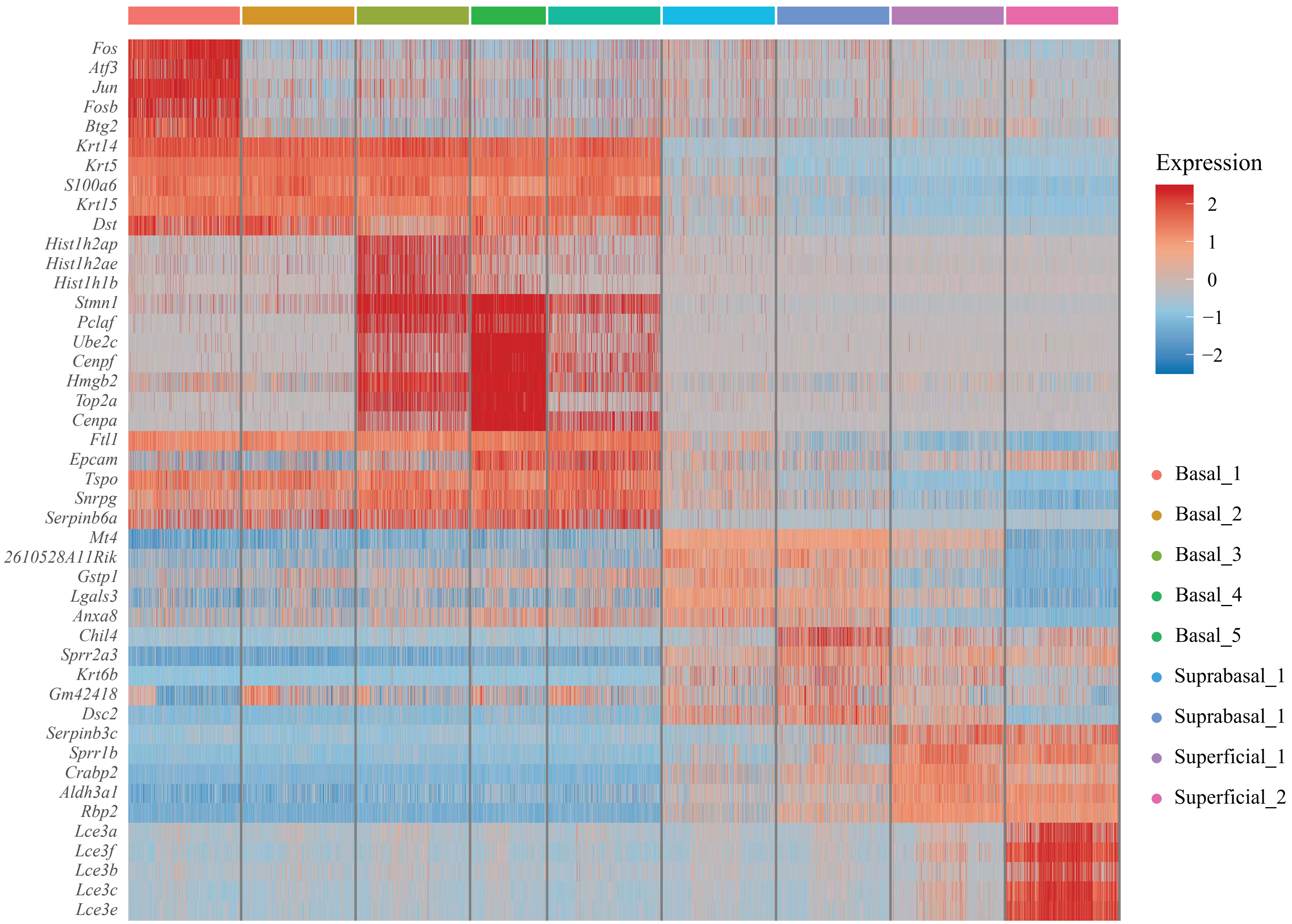

Fos
Atf3
Jun
Fosb
Btg2
Krt14
Krt5
S100a6
Krt15
Dst
Hist1h2ap
Hist1h2ae
Hist1h1b
Stmn1
Pclaf
Ube2c
Cenpf
Hmgb2
Top2a
Cenpa
Ftl1
Epcam
Tspo
Snrpg
Serpinb6a
Mt4
2610528A11Rik
Gstp1
Lgals3
Anxa8
Chil4
Sprr2a3
Krt6b
Gm42418
Dsc2
Serpinb3c
Sprr1b
Crabp2
Aldh3a1
Rbp2
Lce3a
Lce3f
Lce3b
Lce3c
Lce3e
Expression
2
1
0
−1
−2
Basal_1
Basal_2
Basal_3
Basal_4
Basal_5
Suprabasal_1
Suprabasal_1
Superficial_1
Superficial_2


B

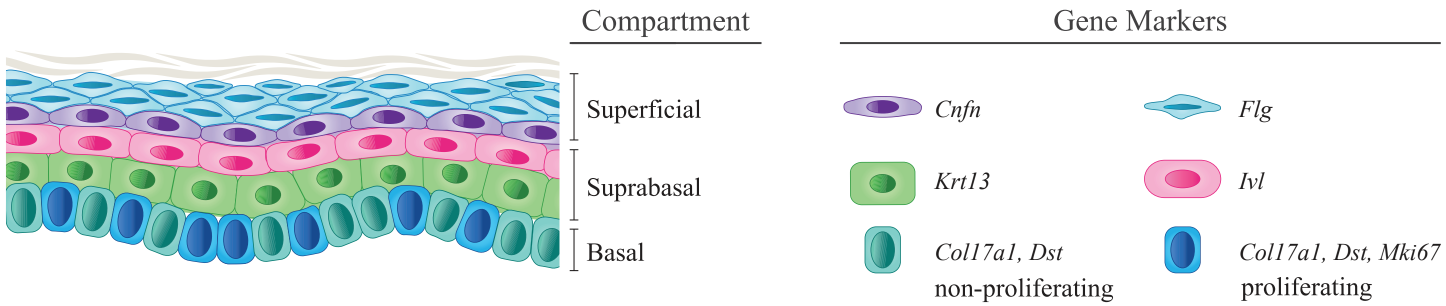

Compartment
Superficial
Suprabasal
Basal
Gene Markers
Cnfn
Flg
Krt13
Ivl
Col17a1, Dst
non-proliferating
Col17a1, Dst, Mki67
proliferating

A

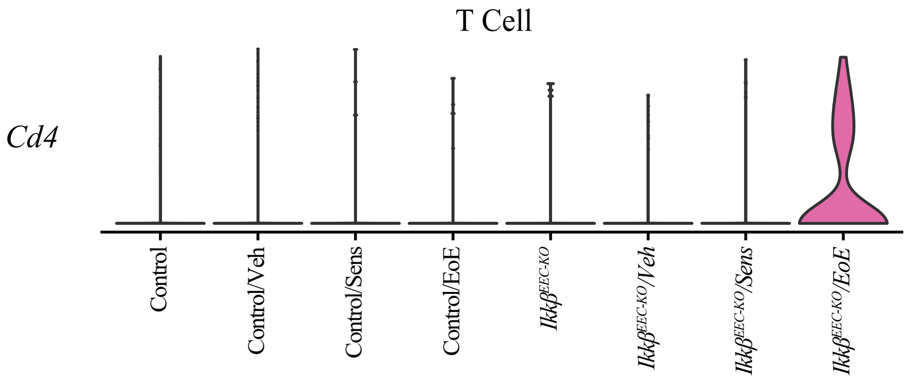
T Cell
*Cd4*
Control
Control/Veh
Control/Sens
Control/EoE
*Ikkβ*EEC-KO
*Ikkβ*EEC-KO/Veh
*Ikkβ*EEC-KO/Sens
*Ikkβ*EEC-KO/EoE

B

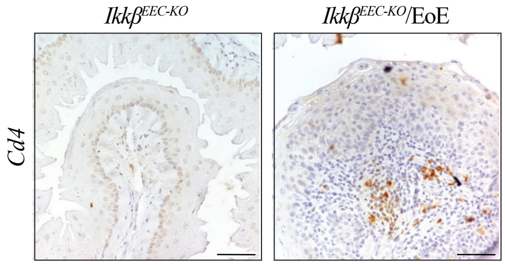
*Ikkβ*EEC-KO
*Ikkβ*EEC-KO/EoE
*Cd4*

**SF8**

A

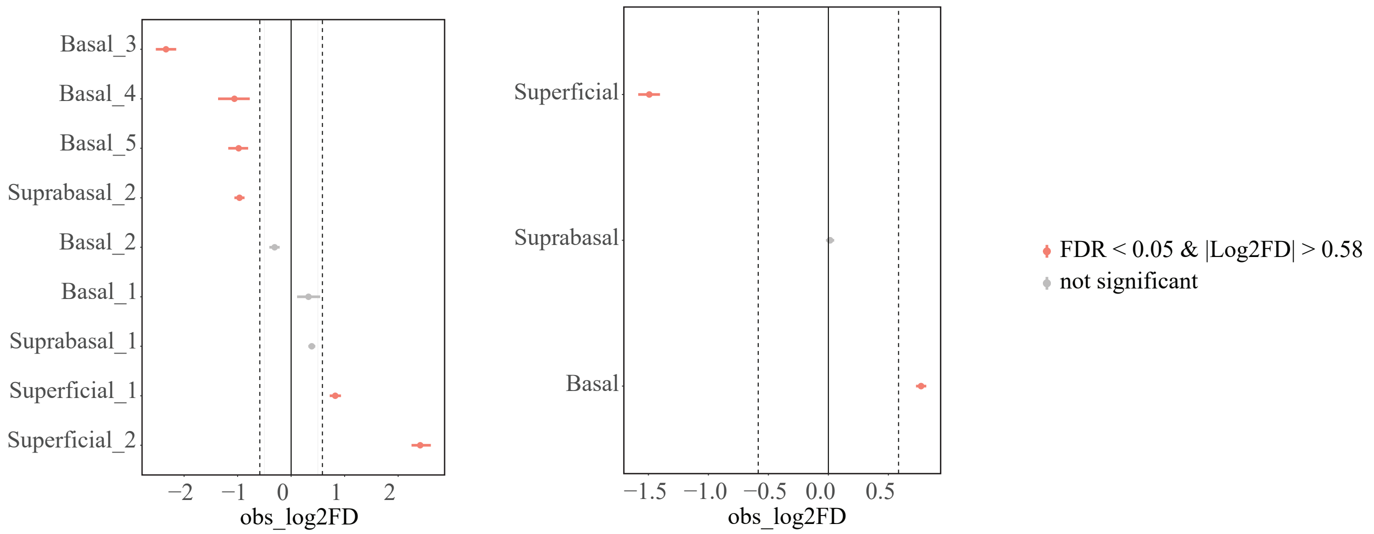
Basal_3
Basal_4
Basal_5
Suprabasal_2
Basal_2
Basal_1
Suprabasal_1
Superficial_1
Superficial_2
−2 −1 0 1 2
obs_log2FD
Superficial
Suprabasal
Basal
−1.5 −1.0 −0.5 0.0 0.5
obs_log2FD
FDR < 0.05 & |Log2FD| > 0.58
not significant

B

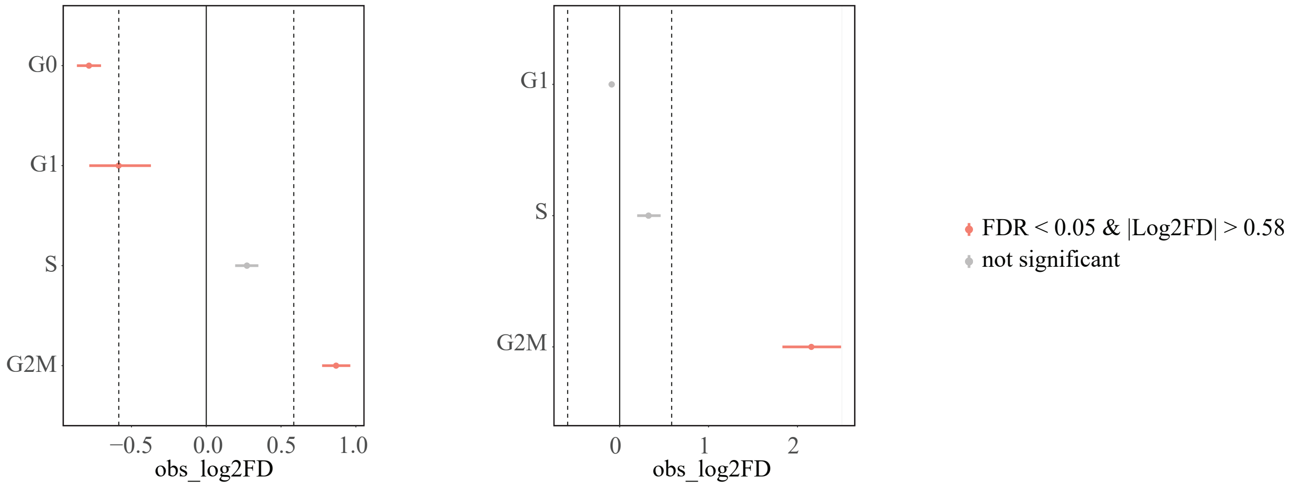
G0
G1
S
G2M
−0.5 0.0 0.5 1.0
obs_log2FD
G1
S
G2M
0 1 2
obs_log2FD
FDR < 0.05 & |Log2FD| > 0.58
not significant

**SF9**

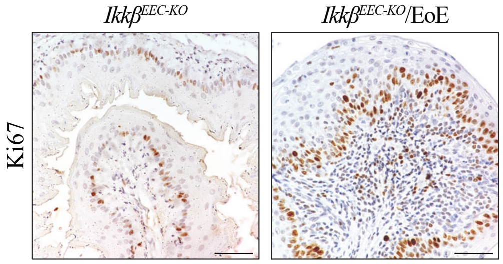

**SF10**

A

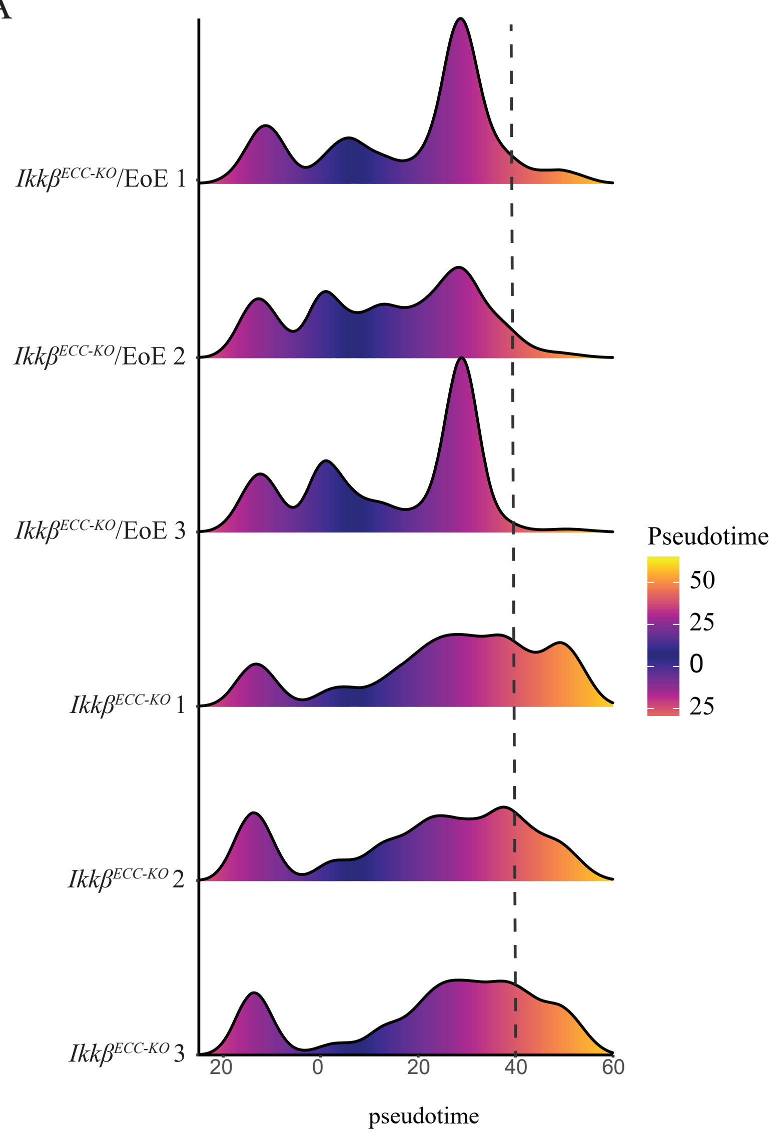
Ikkβ^ECC-KO/EoE 1
Ikkβ^ECC-KO/EoE 2
Ikkβ^ECC-KO/EoE 3
Ikkβ^ECC-KO 1
Ikkβ^ECC-KO 2
Ikkβ^ECC-KO 3
20
0
20
40
60
pseudotime
Pseudotime
50
25
0
25

A

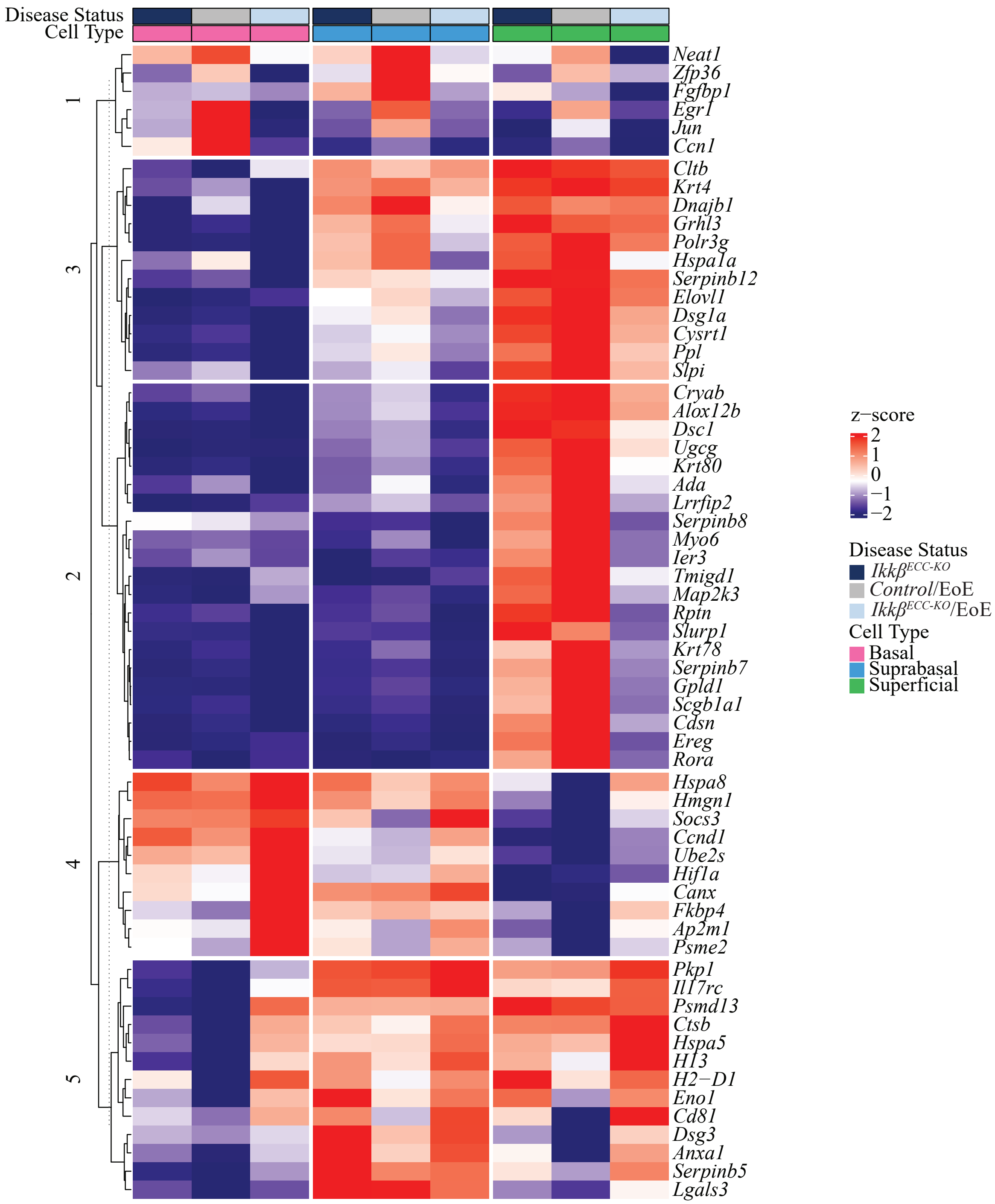
Disease Status
Cell Type
1
3
2
4
5
Neat1
Zfp36
Fgfbp1
Egr1
Jun
Ccn1
Cltb
Krt4
Dnajb1
Grhl3
Polr3g
Hspa1a
Serpinb12
Elovl1
Dsg1a
Cysrt1
Ppl
Slpi
Cryab
Alox12b
Dsc1
Ugcg
Krt80
Ada
Lrrfip2
Serpinb8
Myo6
Ier3
Tmigd1
Map2k3
Rptn
Slurp1
Krt78
Serpinb7
Gpld1
Scgb1a1
Cdsn
Ereg
Rora
Hspa8
Hmgn1
Socs3
Ccnd1
Ube2s
Hif1a
Canx
Fkbp4
Ap2m1
Psme2
Pkp1
Il17rc
Psmd13
Ctsb
Hspa5
H13
H2−D1
Eno1
Cd81
Dsg3
Anxa1
Serpinb5
Lgals3
z−score
2
1
0
−1
−2
Disease Status
Ikkβ ECC-KO
Control/EoE
Ikkβ ECC-KO/EoE
Cell Type
Basal
Suprabasal
Superficial
